# A Unique Type V CRISPR-Cas System Encoded by a Group of *Thermus* Viruses

**DOI:** 10.64898/2026.04.20.719267

**Authors:** Anna B. Trofimova, Alina O. Demkina, Sergey A. Shmakov, Alexei D. Livenskyi, Marina V. Serebryakova, Alexandr A. Dmitriev, Ilya D. Rubinshteyn, Konstantin V. Severinov, Matvey V. Kolesnik

## Abstract

CRISPR-Cas are widespread adaptive immune systems that protect bacteria and archaea from mobile genetic elements such as bacteriophages. Metagenomic sequencing identified CRISPR-Cas systems in phage genomes; however, their functions remain largely unknown. Here, we present Cas12r-CRISPR, a novel type V CRISPR-Cas system encoded by Lalka phages infecting thermophilic *Thermus* bacteria. We determined Cas12r-CRISPR PAM consensus sequence and crRNA structure and showed, that when provided with appropriate spacers and expressed in *Thermus thermophilus*, Cas12r-CRISPR efficiently interferes with plasmid transformation as well as infection by diverse *Thermus* phages. In the course of Lalka phage infection, the Cas12r-CRISPR locus is expressed with middle phage genes and its transcripts are among the most abundant phage RNAs. Notably, most Cas12r-CRISPR spacers target integrative mobile elements widespread in *Thermus* genomes. Both Lalka phages and targeted integrative mobile elements use host tRNA genes as attachment sites. We therefore propose that Cas12r-CRISPR participates in an inter-MGEs conflict.

**GRAPHICAL ABSTRACT:** 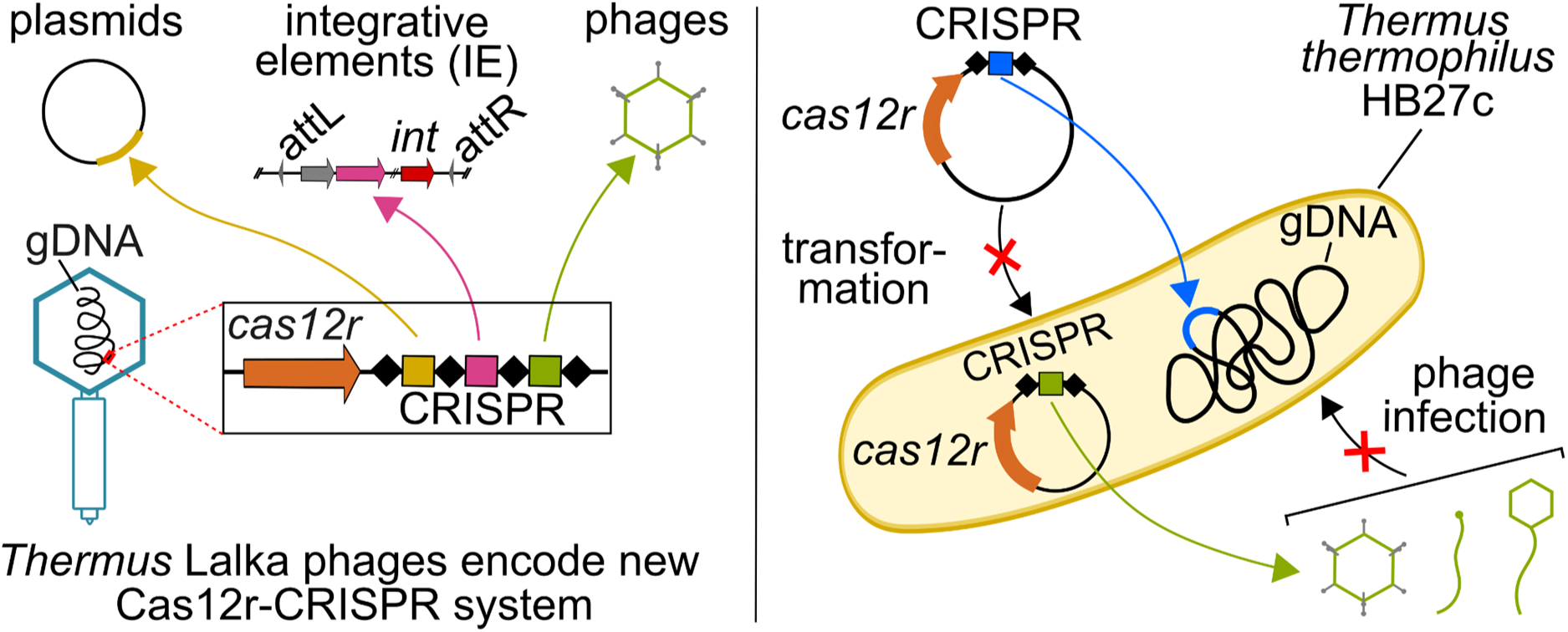

## INTRODUCTION

Viruses and, particularly, bacteriophages are the most ubiquitous biological entities on Earth with viral particles outnumbering cells by an estimated factor of 10 to 100 (1). The antagonistic relationship between bacteriophages and their hosts provides selective pressure that has driven the emergence of a plethora of diverse defense systems in bacteria and, in turn, a variety of counterdefense mechanisms in phages (2). In turn, the excess of viral particles over the host cells drives competition among viruses for the hosts leading to the emergence of mechanisms allowing viruses to thwart the reproduction of the competitors. For example, strictly lytic bacteriophages T4 and T5, encode superinfection exclusion mechanisms that prevent infection by the same or closely related phages (3, 4). Numerous prophages and phage satellites integrated in host genomes encode defense systems (5–7). Such defense systems can be beneficial both for mobile genetic elements (MGEs) and the host cell, and the evolution of MGE-encoded defense systems at least in some cases may be driven primarily by inter-MGE conflict (8). CRISPR–Cas systems, although rare, have also been detected in viral genomes, and analysis of their spacer repertoires suggests that they are involved in interviral conflicts (9–11). Currently, all but one described viral CRISPR–Cas systems have been detected in metagenomic sequences, making it impossible to investigate their activity under native conditions.

The only bacteriophage-encoded CRISPR-Cas system whose biological role has been experimentally demonstrated is a Type I-F CRISPR-Cas encoded by *Vibrio cholerae* phage ICP1 (9). This phage-borne system comprises complete interference and adaptation modules and two CRISPR arrays. The spacers predominantly target PLEs (PICI-like elements), a dedicated family of ICP1 satellites integrated into *V. cholerae* genomes. Upon ICP1 infection, the PLE excises from the chromosome, replicates, and hijacks the ICP1 structural proteins to form viral particles loaded with the PLE genome, which completely prevents the production of the ICP1 progeny (12). The CRISPR-Cas system encoded by some ICP1 lineages allows to restore the reproduction of the phage in the presence of PLEs by acquiring PLE-derived spacers that enable the I-F effector proteins to recognize and destroy the PLE DNA (9). Besides its peculiar biological role which can be described as an antidefense (8), the ICP1 CRISPR-Cas system is remarkable for its unusual mode of action, i.e., recruitment of the Cas1 adaptation protein in the immune response (13).

In this work, we investigated the transcriptional landscape of the *Thermus thermophilus* bacteriophage Lalka27a previously isolated from the Kunashir Island hot springs (14). One of the most abundant Lalka27a transcripts encodes a TnpB-like nuclease followed by a short CRISPR array. We show that this TnpB-like nuclease and the adjacent array comprise a functional CRISPR-Cas system capable of interfering with plasmid transformation or phage infection. Phylogenetic analysis suggests that effectors of CRISPR-Cas systems encoded by Lalka27a and its close relatives define a distinct subtype of Type V systems, which we name Cas12r. To our knowledge, Cas12r-CRISPR is the first Type V CRISPR-Cas system encoded by the viruses of hyperthermophilic bacteria. Spacers from Lalka phages CRISPR arrays match sequences of integrative mobile elements widespread in *Thermus* genomes suggesting a conflict between these elements and Lalka bacteriophages.

## MATERIAL AND METHODS

### Strains and cultivation conditions

*Thermus thermophilus* HB27c (GenBank: GCA_014100815.1) was cultivated in TBM medium (0.8% w/v tryptone, 0.4% w/v NaCl, and 0.2% w/v yeast extract dissolved in ‘Vittel’ mineral water), supplemented with 1 mM MgSO_4_ and 1 mM CaCl_2_ at 70 ℃ in an orbital shaker at 180 rpm. For cultivation on solid medium, 0.8% (w/v) Gelzan CM (Sigma-Aldrich) was added to the TBM medium supplemented with 7 mM CaCl_2_ and 1 mM MgSO_4_ in total. For phage propagation using the double-layer agar method, TBM medium was supplemented with 2% (w/v) agar for the bottom layer and 0.5% (w/v) agar for the top layer. Plates were incubated at 70 ℃ in plastic bags to prevent water evaporation.

For molecular cloning, *Escherichia coli* DH5α (F^-^ *U80lacZDM15* D (*lacZYAargF*) U169 *recA1 endA1 hsdR17*(r_k_^-^, m_k_ ^+^) *phoA supE44 thi-1 gyrA96 relA1* λ^-^) was used. Cells were cultivated in LB medium in a shaker at 37 ℃, 180 rpm. For cultivation on solid media, agar was added to a final concentration of 1.5% (w/v).

*Thermus* bacteriophages phiKo (GenBank: MH673671.2), phiFa (GenBank: MH673672.2), Zuza27 (GenBank: PQ109074), Lalka8 (GenBank: PQ109075), Lalka27a (GenBank: PQ109076), Lalka27b (GenBank: PQ109077) were isolated in our laboratory (14, 15). Phage lysates were prepared by adding a single phage plaque to 10–50 mL of fresh *T. thermophilus* HB27c culture (OD_600_ ∼ 0.2) and incubating for 3–4 h as described above. Cell debris was removed by centrifugation at 21,000 × g for 30 min, and the clarified supernatant was collected and stored at 4 ℃.

### Engineering of plasmids bearing mini-CRISPR arrays

The *lalcas12r* ORF from Lalka8 phage along with its putative promoter region and two CRISPR repeat units was cloned into the pUC19 vector. In the resulting vector, the *lalcas12r* gene is followed by the native 15-bp intergenic region and by two repeat units separated with the stub encoding *lacZα* reporter region. The stub sequence is flanked by Esp3I recognition sites, allowing replacement of the stub with a sequence of interest. The plasmid was built using the Golden Gate assembly approach. BsaI recognition sites were introduced by PCR using primers carrying the required sequence. The PCR fragments were digested with the BsaI restriction enzyme (Thermo Fisher Scientific) and ligated with T4 DNA ligase (Thermo Fisher Scientific). The sequence of the resulting plasmid was validated by the long-read sequencing using Oxford Nanopore MiniION platform. Spacers of interest, flanked by sticky ends complementary to the sequences generated by Esp3I digestion, were prepared by annealing two complementary oligonucleotides (Evrogen, Russia) and cloned into pUC19_*lalcas12r*_mini-CRISPR via Golden Gate assembly, replacing the *lacZα* reporter. Clones containing plasmid with the replacement were selected by blue-white screening. The resulting *lalcas12r*_mini-CRISPR locus, flanked by SalI recognition sites, was excised from the pUC19_*lalcas12r*_mini-CRISPR plasmid using SalI restriction enzyme (Thermo Fisher Scientific) and ligated into the corresponding site of the pMK18 plasmid. Using colony PCR, clones carrying *lalcas12r*_mini-CRISPR locus under the constitutive kanamycin promoter were selected for further experiments. A nucleotide point mutation in one of the catalytically active residues (D238A) of the RuvC domain was introduced using overlap extension PCR. The sequences of the oligonucleotides are provided in **Supplementary Table S1.**

### Determination of temporal classes of Lalka27a genes

Overnight culture of *T. thermophilus* HB27c was diluted 1:50 with fresh TBM medium and cells were grown until OD_600_ ∼ 0.2. The resulting cell culture was infected with Lalka27a (MOI ∼ 10). Aliquots (10 mL) were collected before infection (control) and at 5, 10, 20, 40, 50, and 65 min post-infection. Each sample was immediately supplemented with 1 mL of stop solution (10% phenol in ethanol), placed on ice, and centrifuged at 3,600 × g for 10 min. Cell pellets were resuspended in 50 µL of PBS, followed by the addition of 1 mL of Lira reagent (Biolabmix, Russia). The suspensions were mixed by pipetting, incubated at room temperature for 10 min, and centrifuged at 10,000 g for 10 min at 4 ℃ to remove DNA. Total RNA was extracted using the RNA Purification from Lira Reagent Kit (Biolabmix, Russia) according to the manufacturer’s Total RNA purification protocol.

The resulting RNA samples were treated with RNase-free DNase I (Thermo Fisher Scientific) with the following purification with Lira Reagent Kit (Biolabmix, Russia). Libraries were constructed from 1 µg of total RNA using the MGIEasy Fast RNA Library Prep Set (MGI Tech Co., Shenzhen, China) starting from RNA fragmentation (Section 3.2), without mRNA enrichment. First-strand cDNA was synthesized according to the manufacturer’s instructions; second-strand cDNA was generated with the Directional Second Strand Buffer (with dUTP) to retain strand specificity. End-repair/A-tailing was performed as recommended, followed by ligation of MGIEasy PF Adapters. Adapter-ligated DNA was size-selected and cleaned with the provided DNA Clean Beads, and libraries were amplified for 19 PCR cycles. Library quality and size distribution were assessed using the Agilent High Sensitivity D1000 ScreenTape Assay on a 4200 TapeStation System (Agilent Technologies, Santa Clara, CA, USA), and concentrations were determined with the QuDye ssDNA kit (Invitrogen, Thermo Fisher Scientific, USA). Barcoded libraries were pooled, converted to DNA nanoballs according to the manufacturer’s recommendations, and sequenced on a DNBSEQ-G400 in 2×150 bp paired-end mode. The reads corresponding to *T. thermophilus* ribosomal RNAs were removed with the bbduk.sh tool from BBTools package v39.81 (https://bbmap.org/) using the sequences of 16S and 23S rRNA genes from *T. thermophilus* HB27c genome (GCA_014100815.1) with the k=31 and hdist=1 parameters. Next, reads were subjected to quality filtering and adaptor trimming using trimmomatic v0.39 (16) with the following parameters: PE -phred33 ILLUMINACLIP:adapters:2:30:7 LEADING:0 TRAILING:0 SLIDINGWINDOW:4:0 MINLEN:15. The processed reads were mapped to the reference sequences (Lalka27a genome (GenBank ID: PQ109076.1) and *T. thermophilus* HB27c chromosome and megaplasmid (GCA_014100815.1) using bowtie2 v2.3.5.1 (17) with --very-sensitive-local parameter. The quantification of transcript fragments by phage genes was performed using featureCounts function from the Rsubread package v2.18.0 (18) in reversely stranded mode and allowed multiple overlapping of reads with features. TPM (transcripts per million) values were calculated with normalization on a total number of counted reads. Each Lalka27a gene was assigned to one of the temporal classes - pre-early, early, middle and late - according to its transcript abundance within a certain period of infection. The dynamics of transcript abundance was quantified with the help of a Log-Fold Change parameter (LogFC) that was calculated as follows: LogFC_XvsY = log10A(Y) log10A(X), where A(X) and A(Y) are normalized transcript abundances of the gene at time points X and Y post infection. The maximum values of transcript abundances of the pre-early class genes was expected to be within the first 5 min post infection, so their LogFC values obeyed the following criterion: LogFC_5vs10 < 0 and LogFC_10vs20 < log10(1.5) and LogFC_20vs40 < log10(1.5) and LogFC_40vs50 < log10(1.5) and LogFC_50vs65 < log10(1.5). The abundance of early class genes are expected to reach the maximum at 10 min post-infection: LogFC_5vs10 > 0 and LogFC_20vs40 < log10(1.5). The maximum abundances of the middle genes were expected to be at 20-40 min post-infection: LogFC_10vs20 > 0 and LogFC_40vs50 < log10(1.5) and LogFC_40vs65 < 0. The abundances of the late genes were expected to reach maximum after 40 min post-infection: (LogFC_20vs40 > 0 and LogFC_40vs65 > 0) and (LogFC_40vs50 > 0 or LogFC_50vs65 > 0). The abundances of phage transcripts were visualized with the Gviz v1.48.0 package (19).

### Small RNA isolation, library preparation and sequencing (sRNA-seq)

To isolate small RNAs from *T. thermophilus* HB27c cells expressing *lalcas12r*-CRISPR loci, overnight cultures of the cells harboring distinct plasmids were diluted 1:50 with fresh TBM medium and cells were grown until OD_600_ ∼ 1.0. Then, 10 ml of each culture was centrifuged for 10 min at 3600 g at 4 ℃. Cell pellets were resuspended in 50 µL of PBS, followed by the addition of 1 mL of Lira reagent (Biolabmix, Russia). The suspensions were mixed by pipetting, incubated at room temperature for 10 min, and centrifuged at 10,000 g for 10 min at 4 ℃ to remove DNA. Small RNAs were extracted using the RNA Purification from Lira Reagent Kit (Biolabmix, Russia) according to the manufacturer’s microRNA purification protocol. The resulting small RNA samples were treated with RNase-free DNase I (Thermo Fisher Scientific) and subjected to sRNA-seq. The samples of small RNAs from the Lalka27a infected cells were extracted in the same way.

For small RNA sequencing, libraries were prepared from enriched small RNA using the NEBNext® Small RNA Library Prep Set for Illumina (New England Biolabs, USA) according to the manufacturer’s protocol. The protocol was followed up to the 12-cycles of PCR amplification, after which the reaction product was purified using magnetic beads. Quality was assessed using the Agilent High Sensitivity D1000 ScreenTape Assay on a 4200 TapeStation System (Agilent Technologies Inc., USA) and quantified with the Qubit High Sensitivity dsDNA Assay Kit (Invitrogen, Thermo Fisher Scientific, USA). PCR amplicons were proceeded to end repair, MGI-compatible adapter ligation, circularization, and DNB preparation using the MGI Easy PCR-Free Library Prep Set (MGI Tech Co., Shenzhen, China) in accordance with the manufacturer’s recommendations. Barcoded libraries were pooled in equimolar ratios and sequenced on the DNBSEQ-G400 platform (MGI Tech) in 2×150 bp paired-end (PE) mode. Then, reads in pairs were reordered: reads containing P5 (AATGATACGGCGACCACCGAGATCTACAC) and P7 (CAAGCAGAAGACGGCATACGAGAT) adapter sequences were reassigned to forward and reverse reads respectively with the bbduk.sh tool from the BBTools package using k=13 and hdist=1 parameters. Files containing proper forward and reverse reads were prepared using repair.sh and reformat.sh tools from the BBTools package. The adapter sequences were removed using cutadapt v5.2 (20) with the following parameters: cutadapt -m 15:15 -a AGATCGGAAGAGCACACGTCTGAACTCCAGTCAC -g GTTCAGAGTTCTACAGTCCGACGATC -A GATCGTCGGACTGTAGAACTCTGAAC -G GTGACTGGAGTTCAGACGTGTGCTCTTCCGATCT -n 6. The overlapping paired reads were merged using bbmerge.sh tool from the BBTools package with the mininsert=15 parameter. The merged reads were mapped on the reference sequences (Lalka27a genome and *T. thermophilus* HB27c chromosome and megaplasmid using bowtie2 v2.3.5.1 (17) with --very-sensitive-local parameter. The per-nucleotide coverage was calculated with the bedtools genomecov from the BEDTools package v2.27.1 (21). The coverage was visualized with the pyGenomeViz package (https://github.com/moshi4/pyGenomeViz).

### Determination of promoters and terminators of Lalka27a genes

The promoters in *T. thermophilus* HB27c and Lalka27a genomic sequences were predicted using BPROM web server (http://www.softberry.com/berry.phtml?topic=bprom&group=programs&subgroup=gfindb).

The coverage of 5’ ends of mapped sRNA-Seq reads was assessed manually with the IGV browser (22). To identify promoters which were not predicted by BPROM, intergenic regions of the Lalka27a genome were extracted and analyzed using MEME (23) (the number of motifs: 3, search in a given strand only). The obtained motif was searched in the Lalka27a genome using the FIMO tool (24). Rho-independent transcriptional terminators were predicted using TransTermHP v2.09 (25).

### Phage isolation

Lalka27c and Lalka27d phages were isolated from environmental samples using a previously described approach (14). Aliquots of 5 mL TBM medium were inoculated with 100 µL of the overnight *T. thermophilus* HB27c culture and incubated at 70 ℃ until the OD600 reached ∼0.4. Then, 0.5 mL of environmental sample was added, and the incubation was continued overnight. To isolate individual phage plaques, 1 mL of the enrichment culture was centrifuged at 15,000 g for 15 min at 4 ℃, and 100 µL aliquots of the supernatant were added to 15 mL of melted 0.5% TBM agar supplemented with 200 µL of freshly grown *T. thermophilus* HB27c culture (OD600 ∼0.4). Mixtures were shaken to ensure homogeneity, poured over 2% TB agar plates, and incubated overnight at 70 ℃. Individual plaques were picked using pipette tips and purified by several passages on the host strain.

For phage propagation, host cultures grown until OD_600_∼0.2 were inoculated with a single phage plaque and incubated for 3-4 h. Cultures were centrifuged at 15,000 g for 30 min at 4 ℃. Viral particles were precipitated by overnight incubation on ice with 10% PEG-8000 and 1 M NaCl, followed by centrifugation at 15,000 g for 30 min at 4 ℃. The pellets were resuspended in 500 µL of SM buffer (50 mM Tris-HCl, pH 7.5, 100 mM NaCl, 8 mM MgSO₄).

### DNA extraction from virions

In total, 450 µL of phage lysates or concentrates were treated with 1 µL of RNase A and DNase I (10 mg/mL each) and incubated at 30 ℃ for 1 h. Next, 50 µL of buffer (5% SDS, 100 Tris, 10 mM EDTA, pH 8.0) and 20 µL of proteinase K (20 mg/mL) were added followed by incubation at 56 ℃ for 1 h. DNA was extracted using the phenol–chloroform extraction method, precipitated with ethanol, and resuspended in nuclease-free water.

### Analysis of LalCas12r-associated CRISPR spacers

Spacers from CRISPR arrays associated with LalCas12r were extracted manually. Genomic sequences of 401 Thermacea bacteria were downloaded from NCBI GenBank. Spacers were searched against the Thermaceae genomic sequences using blastn tool with the following parameters: -maxtargetseqs 100 -dust no -word_size 8. Spacer hits along with genomic annotations were visualized using the pgviz package. To identify the PAM motif, only spacer–protospacer alignments without gaps were selected. Alignments with spacer coverage below 0.9 were discarded. For each spacer, 10 bp upstream regions at the 5′ end were extracted. The weblogo was built using the logomaker package (26).

### Plasmid and phage interference assay

Plasmid interference assays were performed in *Thermus thermophilus* HB27c. Cells were transformed with an empty pMK18 vector and pMK18-based plasmids encoding LalCas12r/dLalCas12r and an artificial mini-CRISPR array carrying genome-targeting, self-targeting, or non-targeting spacers according to the protocol described previously (**Supplementary Table S2**) (27). 10 µL of 10-fold serial dilutions of the transformation mixtures were spotted onto TBM-Gelzan plates supplemented with kanamycin (30 μg/ml), and the plates were incubated for 18–42 h at 70 ℃.

For phage interference assays, 10 µL of 10-fold serial dilutions of phage lysates were spotted onto the surface of double-layer TBM agar plates containing kanamycin (30 μg/mL) and freshly seeded lawns of *T. thermophilus* HB27c strains harboring pMK18-based phage-targeting or non-targeting plasmids (**Supplementary Table S2**). Plates were incubated for 18–42 h at 70 ℃.

### Analysis of phiKo phage escapers

An individual plaque of a phiKo phage escaper obtained in the phage interference experiment was added to 10 mL of *T. thermophilus* HB27c culture (OD600 ∼0.2) carrying pMK18_*lalcas12r* with a mini-CRISPR array containing a phiKo-targeting spacer. The cultures (including replicates) were incubated at 70 ℃ with vigorous shaking until complete lysis. Phage genomic DNA was extracted as described above (DNA extraction from virions). The protospacer-containing genomic regions were amplified using primers phiKo_ps_F and phiKo_ps_R, and the resulting amplicons were sequenced by the Sanger method (Center for Collective Use “GENOM”). (**Supplementary Table S1**).

### High throughput sequencing of plasmids and phage genomic DNA

DNA libraries were prepared from 400 ng of plasmid or phage DNA using the MGI Easy PCR-Free Library Prep Set (MGI Tech Co., Shenzhen, China), following the manufacturer’s protocol. The libraries were size-selected and purified using the supplied DNA Clean Beads. Quality control was performed using the Agilent High Sensitivity D1000 ScreenTape Assay on a 4200 TapeStation System (Agilent Technologies Inc., Santa Clara, CA, USA) and quantified with the Qubit ssDNA Assay Kit (Invitrogen, Thermo Fisher Scientific, USA). Equimolar amounts of barcoded libraries were pooled and sequenced on the DNBSEQ-G400 platform (MGI Tech) in 2×150 bp paired-end (PE) mode. Adapters and low quality reads were removed using the fastp tool (28). Plasmid sequences were assembled de novo from the paired reads with unicycler v0.5.1 (29) and annotated with prokka pipeline (30).

### Lalka27a phage purification and total protein extraction

To precipitate viral particles, 150 ml of Lalka27a cell lysate containing ∼ 10^10^ PFU/mL were centrifuged in SW 32 Ti rotor at 30,000 rpm at 4 ℃ for 4 hours. Supernatant was discarded, and the pellet was resuspended in SMC buffer (50 mM Tris, pH=7.5, 200 mM NaCl, 10 mM MgCl_2_, 10 mM CaCl_2_). The obtained sample was layered on the stepwise sucrose gradient (10-60% sucrose in SMC buffer). Centrifugation was performed in a SW 32 Ti rotor at 30,000 rpm for 6 hours at 10 ℃. The resulting sucrose gradient was fractionated into separate samples, to which trichloroacetic acid (TCA) was added to a final concentration of 10%, and incubated overnight at 4 ℃. After centrifugation at 15,000 g for 20 min at 4 ℃, the pellets were resuspended in TCA to remove residual sucrose. Following additional centrifugation, pellets were washed with 50% acetone. The precipitated proteins were dissolved in the Laemmli buffer and separated using SDS-PAGE.

### In-Gel Trypsin Digestion and MALDI-MS Analysis

Pieces (∼1 mm³) of the stained protein-containing SDS–PAGE gel were destained twice with a solution containing 50 mM NH₄HCO₃ and 50% aqueous acetonitrile. Gel pieces were dehydrated with 50 µL of 100% acetonitrile, and rehydrated with 5 µL of the digestion solution containing 15 µg/mL sequencing grade trypsin (Promega) in 100 mM NH_4_HCO_3_. Digestion was carried out at 37°C for 4 h. The resulting peptides were extracted with 5 µL of a solution containing 0.5% trifluoroacetic acid (TFA) and 30% acetonitrile. An aliquot of 1 µL of in-gel tryptic digest extract was mixed with 0.5 µL of 2,5-dihydroxybenzoic acid solution (40 mg/mL in 30% acetonitrile, 0.5% TFA). MALDI-TOF (matrix-assisted laser desorption ionization-time of flight) MS analysis was performed on an UltrafleXtreme MALDI-TOFTOF mass spectrometer (Bruker Daltonics, Bremen, Germany). The MH+ molecular ions were measured in a reflector mode; the accuracy of monoisotopic mass peak measurement was within 50 ppm. Protein identification was carried out using MS ion search with the use of Mascot software version 2.3.02 (Matrix Science) through the home database.

### Phylogenetic analyses

To reconstruct the tree for the Cas12 and TnpB proteins, the approach described previously was used (31). Briefly, Cas12 and TnpB sequences were downloaded from (32) and combined with the sequences described in this paper (Cas12r proteins from Lalka phages and Ga0079992_140590). The following approach was then used: (1) sequences were clustered using MMseqs2 (33) (similarity and coverage thresholds of 0.9); (2) sequences were aligned within each cluster using MUSCLE5 (34); (3) alignment columns were selected with a homogeneity value ≥0.05 and a gap fraction <0.667; and (4) for the resulting alignments, maximum-likelihood trees was constructed using FastTree (35) with the Whelan and Goldman evolutionary model and gamma-distributed site rates.

To determine the hierarchy of poorly alignable clusters, the HHsearch (36) was employed to calculate similarity scores between alignments, followed by the unweighted pair group method with arithmetic mean (UPGMA) approach, as described in (31).

To reconstruct the tree for the RuvC domains of the Cas12 and Type V-U families, RuvC profiles from (31) were used for the previously described CRISPR-associated TnpB-like protein families. For the sequences presented in this study (Cas12r proteins from Lalka phages and Ga0079992_140590), MUSCLE5 (34) was used with the ‘super5’ option to align the sequences. To identify the RuvC domains within the alignment, new sequences were aligned against known RuvC domains. Then the pipeline presented above was employed, as detailed in (31), to construct the final tree of the RuvC domains.

### Evaluation of CRISPR-Cas systems diversity in *Thermus* bacteria

CRISPR-Cas systems diversity was evaluated with PADLOC v2.0.0 (37). Open reading frames were predicted using Prodigal v2.6.3 (38) in 224 *Thermus* genome sequences downloaded from the NCBI database. The resulting predictions were used as an input for the PADLOC pipeline with “--fix-prodigal” option. Two predicted Type V loci were removed as false positives after manual revision due to the lack of adjacent CRISPR arrays. Standalone cas_adaptation modules predicted by PADLOC were also discarded. The results of the PADLOC pipeline are provided in **Supplementary Data File S1**.

### Detection of tRNA-specific integrative elements in *Themus* genomes

To estimate the abundance of integrative elements (IEs) that use tRNA genes as attachment sites in *Thermus* bacteria, we developed a pipeline called MGE_finder (https://github.com/rljech13/MGE_finder). Protein-coding genes were predicted with Prodigal v2.6.3 in single-genome mode, yielding amino acid sequences and coding sequence coordinates. Tyrosine integrase genes were searched using hmmscan with HMM profiles PF00589 and PF22022 from the Pfam database (39), yielding a list of *Thermus* integrase hits. The location of tRNA genes in the *Thermus* genomes were predicted with ARAGORN v1.2.41 (40). Integrase genes located on the opposite strand within 500 bp downstream of the nearest tRNA gene were considered as tRNA-associated. Nucleotide sequences of the integrase-adjacent tRNAs were used as a query for blastn v2.16.0 (41) to search for homologous sequences within 100 kbps upstream the integrase gene. Predicted hits were filtered to identify candidate attachment site organizations with an intact tRNA gene located downstream of the integrase gene and a tRNA 3′ segment sequence located upstream of the integrase gene. The pipeline was implemented using Snakemake v7.32.4 (42).

## RESULTS

### Transcription strategy of the Lalka27a phage

To investigate transcription of bacteriophage Lalka27a genes throughout the infection, a culture of *T. thermophilus* HB27c cells was synchronously infected with the phage at high multiplicity of infection (MOI ∼ 10) and culture aliquots were collected 5, 10, 20, 40, 50, and 65 minutes post-infection (at our conditions, the release of phage progeny аnd cell lysis start 70 minutes post-infection). Total RNA was isolated from cells collected at each time point, subjected to RNA-Seq, and reads were mapped on the host and phage genomic sequences. Phage transcripts started to appear at the earliest time-point and, by the end of the infection, constituted 36% of total RNA reads (**Supplementary Fig. S2A**). The results of phage reads mapping are presented in **Fig. 1**. Since the true structure of Lalka27a genome ends is not known, the genome is presented in a state corresponding to a Lalka-like prophage observed in the *Thermus sp.* 93170 genome (NZ_JBPXNL010000001.1:1975845-2048918) (**Fig. 1A**, **Supplementary Fig. S1**). Based on the accumulation kinetics of phage transcripts throughout the infection, Lalka27a genes were divided into immediate early (pre-early), early, middle, and late temporal classes (**Fig. 1A**, **Supplementary Fig. S2A, S2B**, and **Supplementary Table S3**).

**Figure 1.**
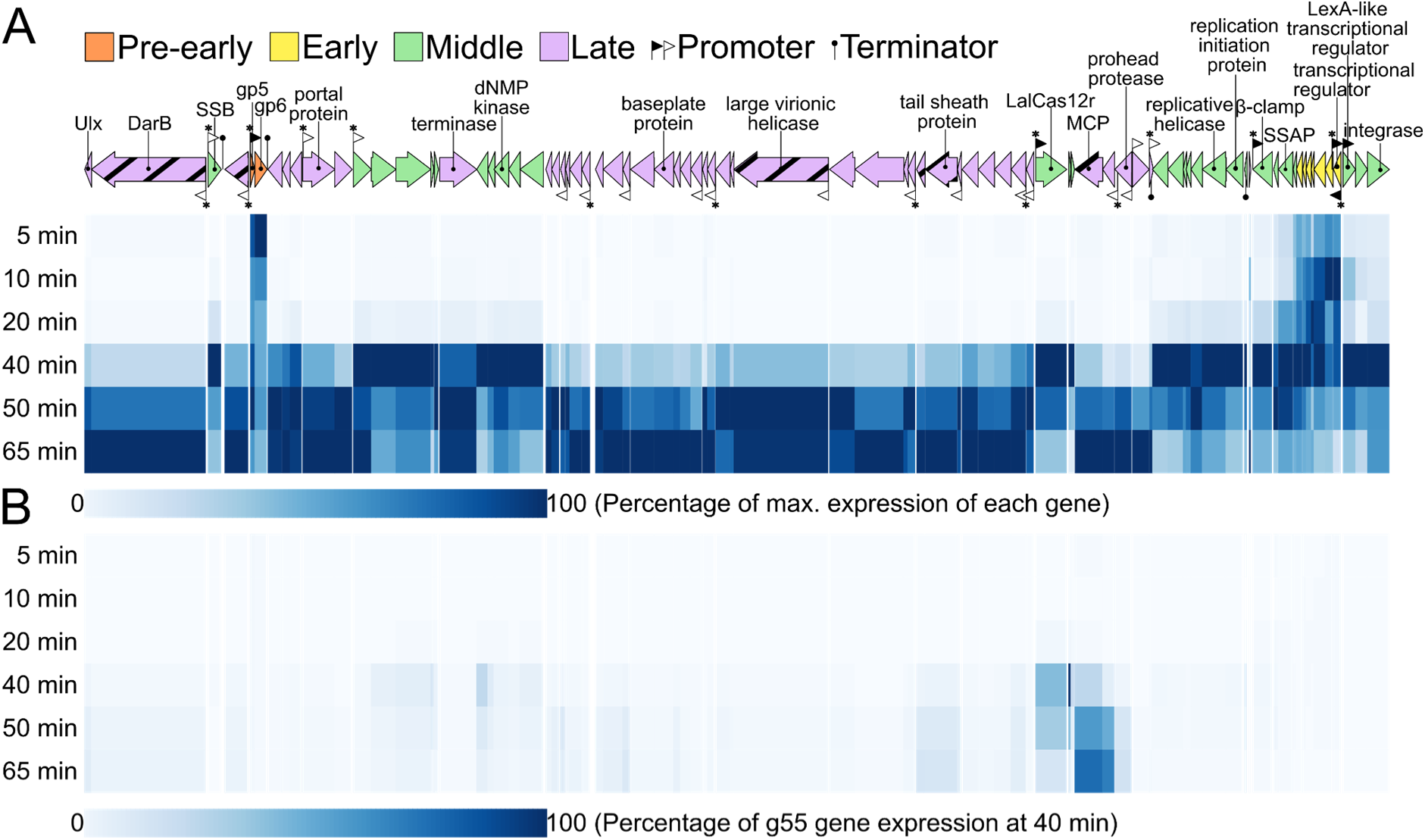
Analysis of Lalka27a genes transcription during infection. At the top, the Lalka27a genome is schematically presented. Open reading frames are shown as arrows. Genes encoding proteins identified in Lalka27a virions are shown as hatched arrows. Promoters predicted by the BPROM tool are indicated with black filled flags. Promoters predicted through motif enrichment analysis are indicated with white flags. Asterisks indicate promoters supported by sRNA-Seq data. Predicted transcription terminators are indicated with black filled circles. Heatmaps indicating abundance of Lalka27a transcripts throughout the infection are shown below. (**A**) Transcript abundance for each gene normalized to the maximal abundance of this gene throughout the infection. (**B**) Transcript abundances normalized to the highest absolute abundance attained by a phage transcript (gene *55* at 40 min post-infection).

The transcript abundance of two pre-early Lalka27a genes, *5* and *6*, is highest 5 minutes post-infection. Gp5 is a small (82 amino acids, ∼10 kDa) acidic (predicted pI 4.41) protein; Gp6 is a medium-sized (222 aa, ∼25 kDa) protein with predicted pI of 4.94. A close homolog of Lalka27a Gp5 (84% of identity of amino acid sequence) is encoded by *Thermus* myovirus phiMN1 (43). The Lalka phages share only ten homologous genes with phiMN1, and the Gp5 homologs pair is by far the closest one (**Supplementary Fig. S3**). The high conservation of the amino acid sequences suggests that Gp5-like proteins play an important role in both phages. Similar to other acidic phage antirestriction proteins, Lalka27a Gp5 and Gp6 may suppress host defense systems targeting foreign nucleic acids by acting as nucleic acid decoys (44, 45).

The early Lalka27a genes (*75–81*) are organized in a leftward-transcribed cluster and likely form a single operon (**Fig. 1A**). Their transcripts are detectable 5 minutes post-infection and reach maximal abundance at later times. Homology modeling allows to predict the function of only one early protein, Gp81, a putative helix-turn-helix transcription factor

Several clusters of Lalka27a middle genes are scattered across the genome (**Fig. 1A**). The middle genes cluster located at the right end of the genome is transcribed rightward and comprises genes encoding a LexA-like transcriptional regulator, a protein with a predicted transmembrane domain, and an integrase that is likely responsible for integration in the host genome. Other middle genes encode proteins involved in DNA replication and nucleotide metabolism: a single-strand annealing protein (SSAP), a β-clamp, a replicative helicase, a dNMP kinase, and a single-strand binding protein (SSB). The most abundant Lalka27a middle transcript encodes a predicted TnpB-like nuclease (**Fig. 1B**). In Lalka27a and in closely related Lalka8 and Lalka27b phage genomes, genes encoding this protein are followed by CRISPR-like arrays comprising several 29-bp degenerate repeats separated by unique spacers. As it was previously suggested that TnpB proteins have been recruited as type V CRISPR–Cas effectors on multiple independent occasions (32), we hypothesized that Lalka-encoded TnpB proteins may function as type V CRISPR-Cas effectors (14). Hereafter, we refer to Lalka-encoded TnpB proteins as LalCas12r.

As expected, many late genes in the Lalka27a genome encode predicted virion proteins. A homolog of the *Escherichia coli* P1 phage virion protein DarB (Gp2 in Lalka27a) is also encoded by one of the late genes. The protein composition of sucrose gradient–purified Lalka27a virions was determined by SDS-PAGE analysis followed by tryptic peptide mass fingerprinting. The list of Lalka27a virion proteins is presented in **Supplementary Table S4**. As expected, Lalka27a-encoded Gp2 is a minor component of the Lalka27a virion, supporting a proposed antidefense role like that of phage P1 DarB (46). Another large (1715 aa) virion-associated protein gene is located within a cluster of structural genes. This protein, further referred to as large virionic helicase (Lvh) contains a C-terminal SF2 superfamily helicase domain and is presumably injected into cells during infection. Previously, a number of large multidomain proteins (polyvalent proteins) were shown to be widespread among phages and conjugative plasmids and were predicted to be injected into cells during early stages of infection or conjugation, where they may possess anti-defense functions (47).

The Lalka27a phage does not encode recognizable RNA polymerase or RNA polymerase sigma factors and must therefore rely on host machinery for transcription of its genes. Putative *Thermus* RNA polymerase σ^A^ holoenzyme promoters in the Lalka27a genome were predicted using the BPROM tool. From a total of 32 predicted promoters, 26 were located deep inside the coding sequences and/or had an orientation opposite to the expected direction of transcription. The six remaining putative promoters may initiate the synthesis of pre-early, early, and middle phage transcripts (**Fig. 1A**, **Supplementary Fig. S5**).

In addition to standard RNA-Seq, we performed global sequencing of enriched short RNA fractions from infected cells (sRNA-Seq, see also below). Mapping of reads from enriched fractions revealed RNAs whose 5’ ends were located 6 nucleotides downstream of the BPROM-predicted TANNNT motif corresponding to the *Thermus* −10 promoter consensus element (48) in five predicted phage promoters (no sRNA-Seq signal was observed for P82-M, a putative rightward promoter located in front of middle gene *82*). Bacterial RNA polymerases typically initiate transcription at purines located 6 nucleotides downstream of the −10 promoter element (49, 50). To validate the unexpected observation that sRNA-Seq identifies transcription start sites (TSSs), we inspected reads mapping to several randomly chosen promoters predicted by the BPROM tool in the *T. thermophilus* HB27c genome. As is shown in **Supplementary Fig. S4**, in all cases enriched 5’ ends of short RNAs coincided with annotated TSS positions. We conclude that at least in our model system, sRNA-Seq analysis allows one to identify *in vivo* TSS positions.

By considering enriched sRNA-Seq reads whose 5’ ends are located six nucleotides downstream of the TANNNT motif as TSSs, we identified additional promoters of phage genes (P3-M, P12-M, putative rightward promoters located in front of middle genes *3* and *12*, and P61-L, a putative rightward promoter located in front of the late gene *61*) (**Fig. 1A, Supplementary Fig. S5A, S5B**). Given that the latest time point in the sRNA-seq data was 40 min post-infection, some late promoters may have been missed. To identify other late promoters, we used MEME to detect motifs enriched in intergenic regions upstream of late phage genes. In this way, sixteen putative late promoters containing the highly conserved TAGGGT motif were identified (**Supplementary Fig. S5B**). For half of them, TSSs located six base pairs downstream of the −10 promoter element were observed in sRNA-Seq data. For late promoters that lacked sRNA-Seq support, their genomic location was considered to select the most plausible MEME predictions. For example, two oppositely oriented late promoters in the intergenic region separating divergently transcribed late genes *59* (encoding a prohead protease) and *60* (encoding a recombinase) are most certainly correctly predicted by MEME, as no other sequences that can serve as promoters are present in this intergenic regions.

Although all phage promoters contain a TANNNT motif, the consensus sequences differ between promoters of different temporal expression classes. The combined logo of one pre-early, one early and six middle promoters matches that of host σ^A^ promoters, with a −35 TTGACA consensus and the −10 TAxxxT consensus (in some cases, the −10 promoter element is extended with an upstream TGx motif and the −35 element is lacking) (**Supplementary Fig. S5A**). The only promoter from this group that does not confirm to the consensus is P12-M, which appears to lack a recognizable −35 element and has a −10 element without the TGx extension (**Supplementary Fig. S5A**). All but one late promoter have a distinct TAGGGT −10 element with an additional universally conserved cytosine at position - 25 (**Supplementary Fig. S5B**). This unique consensus suggests that promoter specificity of host RNA polymerase is modified by a phage encoded transcription factor at the onset of late transcription. As previously observed by us for the *Thermus* phage YS40 (51), most Lalka27a transcripts are leaderless, with the adenine or guanine of the initiator codon located at the 5′ end of the transcript (**Supplementary Fig. S5**).

We were not able to identify the promoter(s) responsible for leftward transcription of the middle *22-18* gene cluster located downstream of a leftward-transcribed cluster of late genes *23-25* (**Fig. 1A**). These middle genes may be transcribed from an intragenic promoter located inside one of the late genes. Several predicted promoters supported by sRNA-Seq reveal details about potential regulation of Lalka27a gene expression. Promoter P71a-M (validated by sRNA-Seq data) produces an antisense transcript to the transcript of the β-clamp gene *71* and may thus regulate phage genome replication. The divergently oriented P81-E and P82-M promoters are located in the intergenic region separating genes *81* and *82*, which encode a putative helix-turn-helix transcriptional regulator and a LexA-like transcriptional regulator, respectively. This organization resembles that of the immunity region of *Escherichia coli* bacteriophage λ: the products of genes *81* and *82* may be functional analogs of λ *cro* and *cI* genes, with P81-E and P82-M corresponding to λ P_R_ and P_RM_ promoters, respectively. The lytic/lysogenic switch of λ phage also includes the P_RE_ promoter located downstream of the *cro* gene; P_RE_-initiated transcription proceeds in a direction opposite to that of the *cro* gene (52). The rightward oriented P80a-M promoter inside Lalka27a gene *80* may therefore have the same function as λ P_RE_. Thus Lalka phages, which are capable of lysogenising its host as evidenced by a Lalka-like prophage in the *Thermus sp.* 93170 genome, may be relying on a control strategy similar to that of phage λ. Presumably, lysogenization occurs through the function of Lalka27a integrase, the product of gene *84* that is transcribed together with the LexA-like transcriptional regulator gene.

Using standard tools, we also predicted several intrinsic transcription terminators separating collinearly transcribed phage genes (**Fig. 1A**). Most predicted terminators are in intergenic regions separating convergently transcribed phage genes and may thus be bidirectional. The high density of late promoters and the lack of obvious terminators separating collinearly transcribed genes suggests that unlike λ, Lalka27a does not rely on antitermination mechanisms to control its transcription.

### Lalka phage-encoded *cas12r*-CRISPR loci contain diverse spacers matching *Thermus* mobile genetic elements

To expand the diversity of *lalcas12r*-CRISPR loci, we isolated and sequenced two additional Lalka-like phages, Lalka27c and Lalka27d, from water samples collected at the Valentina hot spring (Kunashir Island) in July 2025. We also searched for homologs of LalCas12r proteins in NCBI non-redundant proteins, IMG/VR, and MGnify databases (53–55) and identified three metagenomic contigs containing *cas12r*-like genes and adjacent CRISPR arrays (**Supplementary Fig. S6**). At the level of nucleic acid sequences, contigs Ga0181858_1004277 (assembled from compost metagenome) and ERZ4756919.5876 (assembled from Diamante hot spring metagenome) share similarity with Lalka phages suggesting that they represent genome fragments of these bacteriophages or prophages. The Cas12r-like proteins encoded by these two contigs are highly similar (>99% identity) to LalCas12r homologs, and we therefore also refer to them as LalCas12r (**Supplementary Fig. S6A**). The Ga0079992_140590 contig (metagenome of microbial community from Yellowstone hot spring) encodes a distinct variant of a Cas12r-like protein that is ∼67% identical to LalCas12r. We therefore refer to it as LlkCas12r (Lalka-like Cas12r) (**Supplementary Fig. S6B**). Although the *llkcas12r*-CRISPR locus is also located among predicted phage genes, its genomic context is different from that of Lalka phages and it must be carried by a distinct phage.

A total of 26 unique spacers were extracted from five Lalka phage genomes, one Lalka-like prophage from *Thermus sp.* 93170, and two metagenomic contigs containing *lalcas12r*-CRISPR loci. All spacers were associated with degenerate 29-bp repeats that were distinct from the *llkcas12r-*associated repeats (**Supplementary Fig. S7**). While most spacers were 31-33 nucleotides long, the Lalka27b and Lalka27c CRISPR arrays shared an identical six nucleotide spacer (shown as a rectangle with diagonal crosses in **Fig. 2A**), which was excluded from further analysis. Of the remaining 25 unique spacers, 7 did not match any known sequences, whereas 16 had matches in the genomes of various *Thermus* isolates and 2 - in *Thermus* plasmids (**Fig. 2A, Supplementary Table S5**). At least one matching spacer was present in every *lalcas12r* array (**Fig. 2A**). Type V CRISPR-Cas effectors recognize PAMs located proximal to the 5’ ends of matching protospacers (56). A 5’ consensus ARG motif was identified immediately adjacent to sequences that matched LalСas12r-CRISPR spacers (**Fig. 2B**). This motif is a likely PAM recognized by the LalCas12r effector.

**Figure 2.**
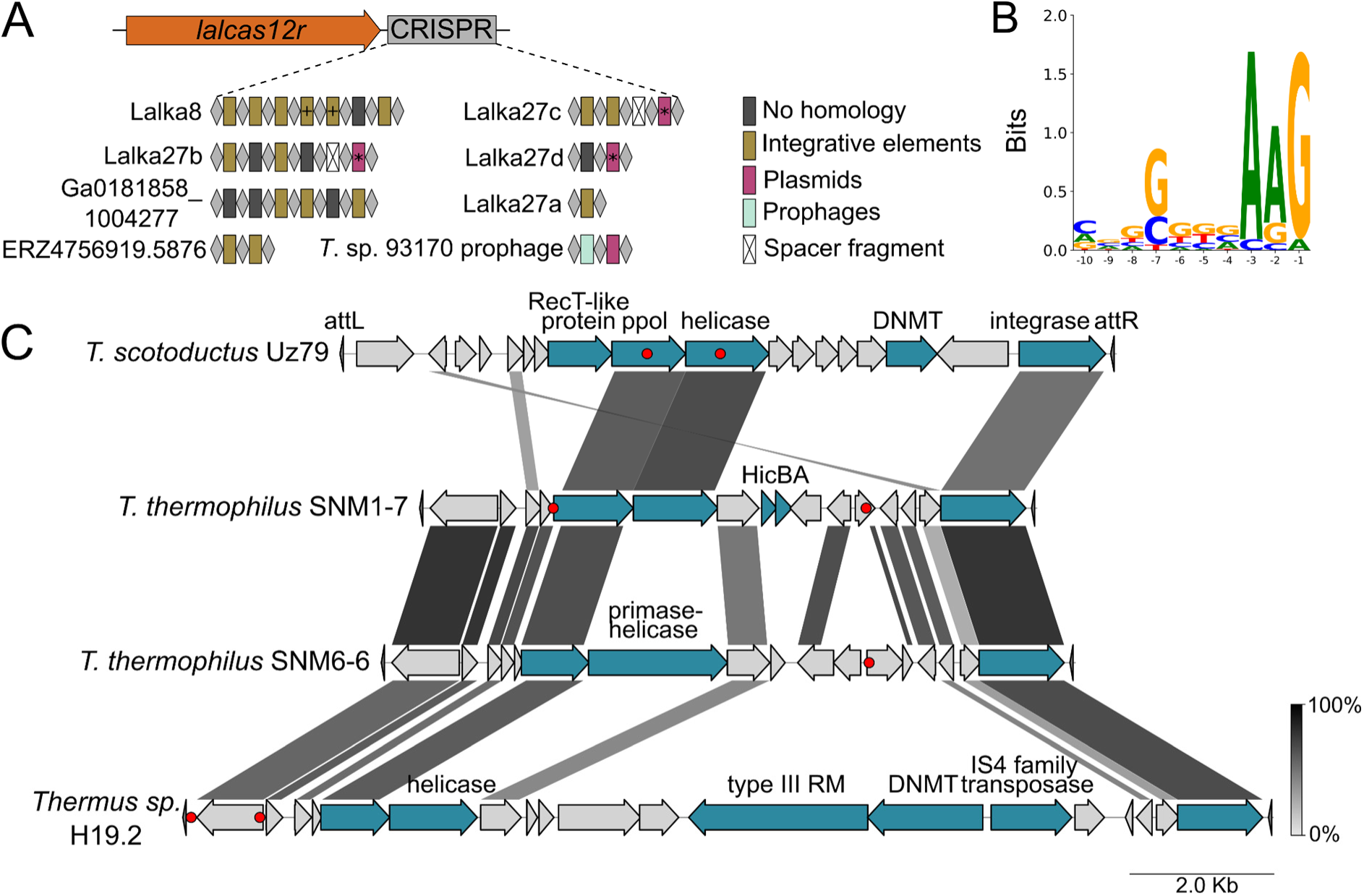
The diversity of spacers encoded by *lalcas12r*-CRISPR loci. (**A**) The organization of the *lalcas12r*-CRISPR locus is shown at the top. Below, the content of CRISPR arrays from 5 Lalka phages, one Lalka-like prophage from *Thermus sp.* 93170, and 2 metagenomic contigs is shown. The repeat units and spacers are depicted, respectively, as grey diamonds and rectangles of different colours. Spacers are colored according to the genomic context of matching protospacers. Spacers without matches are colored dark grey. Spacers with identical sequences are marked with asterisks or crosses. The 6-nt long spacer fragments are marked as white rectangles with diagonal crosses. (**B**) A weblogo of a motif adjacent to the 5’ ends of protospacers matching LalCas12r spacers. (**C**) Graphical alignment of *Thermus* integrative elements (IEs) containing protospacers matching LalCas12r-CRISPR spacers. Homologous gene segments shared between IEs are connected by shading, with shading intensity corresponding to amino acid sequence identity, as shown in the bar at the bottom right. The locations of protospacers matching LalCas12r spacers are shown with red circles.

15 LalCas12r-CRISPR-targeted protospacers were located upstream (within ∼15 kb) of site-specific tyrosine integrase genes present in some *Thermus* isolates (**Supplementary Table S5**). The integrase genes were immediately followed by an intact tRNA gene. In all cases, a 42-46 bp fragment corresponding to the 3′ terminal end of the same tRNA gene was identified within ∼16 kb upstream of the intact tRNA gene. We assume that direct repeats formed by the complete and partial tRNA gene copies arise from site-specific recombination of integrative elements (IEs) catalysed by the product of the integrase gene. All protospacers matching LalCas12r spacers were located between the complete and the partial tRNA gene (four representative cases are presented in **Fig. 2C**). Thus, IEs that use host tRNA genes as attachment sites are apparently targeted by spacers of the LalCas12r-CRISPR system. Although most IE-encoded proteins are unique and of unknown function, all IEs encode proteins containing predicted primase-polymerase (ppol) and helicase domains (**Fig. 2C**). Previously, a related mobile genetic element ICETh2 from *T. thermophilus* HB27 inserting into the tRNA-Val gene was characterized (57). ICETh2 is able to excise from and integrate into host DNA by employing an integrase it encodes. Strikingly, LalCas12r-CRISPR-targeted integrases in *Thermus* IEs are very similar to those of Lalka phages (**Supplementary Fig. S8**). In fact, the Lalka-like prophage in *T. sp.* 93170 (**Supplementary Fig. S1**) is inserted at the tRNA-Ala gene. Thus, Lalka phages and IEs appear to use the same attachment sites.

The three remaining unique LalCas12r spacers match other *Thermus* mobile genetic elements (**Supplementary Table S5**). Specifically, one spacer (present in three different LalCas12r-CRISPR arrays) matches a contig that encodes a *ppol* gene disrupted by an insertion sequence and a gene encoding a TraK-like protein presumably involved in conjugative transfer. The second spacer, encoded by the *T. sp.* 93170 Lalka-like prophage, matches a previously described Mu-like prophage of *T. aquaticus* Y51MC23 (58), while the third one matches a 4.6 kbp contig encoding a protein similar to RepA from *Thermus sp.* ATCC 27737 plasmid (59).

### LalCas12r is a functional CRISPR-Cas system

To test the activity of the Lalka Cas12r-CRISPR system, we prepared a set of plasmids based on the *Thermus*-*E. coli* pMK18 shuttle vector encoding variants of the *lalcas12r*-CRISPR locus. The *lalcas12r* gene was cloned with its 5’ upstream region containing the middle phage promoter. Downstream of the *lalcas12r* open reading frame (ORF), there was a native 15-bp sequence separating the 3’ end of *lalcas12r* and the first CRISPR repeat followed by a spacer and another repeat copy. The plasmids are hereafter referred to as pMK18_*lalcas12r*–r_spacer_r vectors, where “spacer” defines the target.

Although a putative ARG PAM was inferred from sequences adjacent to the 5′ ends of protospacers matching LalCas12r–CRISPR spacers (**Fig. 2B**), LalCas12r–CRISPR-mediated targeting may require additional elements that were not detected due to the limited number of available sequences. To address this, we chose three 8-nt sequences located immediately upstream (5′) of protospacers within *Thermus* IEs that fully matched Lalka spacers (**Supplementary Table S6**). We assumed that the selected sequences contained PAMs that enable targeting. We next searched the *T. thermophilus* genome, as well as the pMK18 plasmid sequence, for exact matches to chosen 8-nt sequences. Two 32-bp sequences located downstream of arbitrary chosen matches in the *Thermus* genome were inserted into the mini-CRISPR arrays of the pMK18_*lalcas12r*–r_spacer_r vector as spacers. The resulting genome-targeting plasmids GT1 and GT2 were expected to be toxic for *T. thermophilus* HB27c cells due to LalCas12r–CRISPR-mediated targeting of the host genome. Another 32-bp target sequence adjacent to putative PAM was selected from the pMK18 plasmid and used to construct a self-targeting (ST) plasmid. This plasmid was expected to be unstable in *T. thermophilus* HB27c cells; thus, no transformants on medium containing kanamycin, which selects for plasmid-carrying cells, should be observed if the Cas12r-CRISPR system is functional.

The GT and ST plasmids, along with empty pMK18 and an NT plasmid were transformed into *T. thermophilus* HB27c cells **(Fig. 3A)**. The latter plasmid contains a spacer for which no matching sequence is present in either the *T. thermophilus* HB27c genome or pMK18. 10-fold serial dilutions of transformation mixtures were plated onto the solid medium supplemented with kanamycin. As can be seen from **Fig. 3B, Supplementary Fig. S9**, the GT and ST plasmids transformed *T. thermophilus* HB27c cells with efficiencies two to three orders of magnitude lower than those of empty pMK18 or NT plasmids. We therefore conclude that the

**Figure 3.**
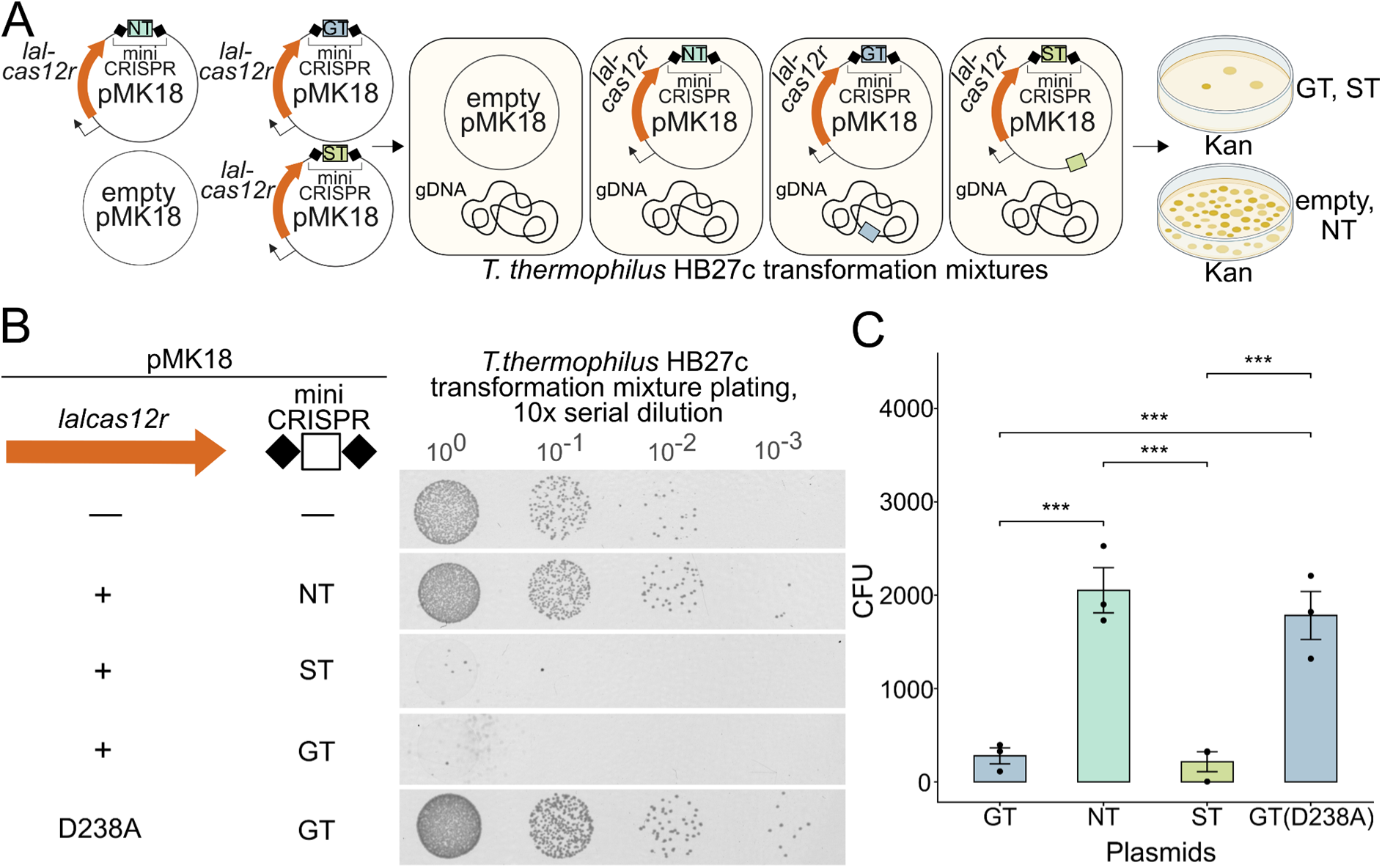
The Lalka-derived Cas12r–CRISPR system interferes with plasmid transformation in *T. thermophilus* HB27c. (**A**) The workflow of plasmid transformation interference assay. *T. thermophilus* HB27c cells were transformed with pMK18 or pMK18-derived plasmids encoding *lalcas12r* and a mini-CRISPR array carrying either genome-targeting (GT), self-targeting (ST), or non-targeting (NT) spacers, followed by plating of transformation mixtures on medium selecting for plasmid-bearing cells. (**B**) The overnight growth of aliquots of 10-fold serial dilutions of indicated transformation mixtures on selective medium. D238A denotes a GT plasmid encoding a catalytically inactive LalCas12r variant (D238A). **(C)** The transformation efficiency of *T. thermophilus* HB27c cells with distinct *lalcas12r*-CRISPR-bearing constructs. Asterisks indicate statistically significant differences (adjusted p-value < 0.001, pairwise t-test with Benjamini–Hochberg correction). Source data are provided in Supplementary Table S7.

Lalka-derived Cas12r-CRISPR system is active and interferes with plasmid transformation in *T. thermophilus* cells. Given the observed lack of transformation of the ST plasmid in *Thermus* and our ability to construct this plasmid in *E. coli*, the LalCas12r–CRISPR system is either not functional or not expressed in *E. coli* cells.

Most of the active Cas12 nucleases, as well as many RNA-guided TnpB nucleases, rely on a conserved RuvC domain to cleave their targets (56). An active RuvC domain contains three conserved negatively charged amino acid residues within the catalytic site (32). Substitution of one of these catalytic residues in LalCas12r, D238, for an alanine abolished interference with GT plasmid transformation, demonstrating that the activity of Lalka Cas12r–CRISPR depends on the RuvC domain (**Fig. 3B**).

Next, we assessed the activity of LalCas12r-CRISPR against *Thermus* bacteriophages. We selected targets within the genomes of three distinct *Thermus* bacteriophages - phiKo (*Tectiviridae*), Zuza27 (*Inoviridae*), and phiFa (*Siphoviridae*) (14, 15). Spacers matching targets adjacent to sequences corresponding to PAMs validated in the plasmid transformation assay (above) were inserted into mini-CRISPR arrays of *lalcas12r*-CRISPR loci encoded on pMK18-based vectors, resulting in phage targeting (PT) plasmids. These plasmids, along with the NT plasmid, were introduced into *T. thermophilus* HB27c cells, and the resulting cultures were challenged with bacteriophages in plaque-forming assays. As can be seen from **Fig. 4**, the targeting of the phage genome with LalCas12r-CRISPR reduced the number of plaques by 4–5 orders of magnitude compared to the NT control. Thus, the Lalka-derived Cas12r–CRISPR system efficiently interferes with infection by diverse *Thermus* phages.

**Figure 4.**
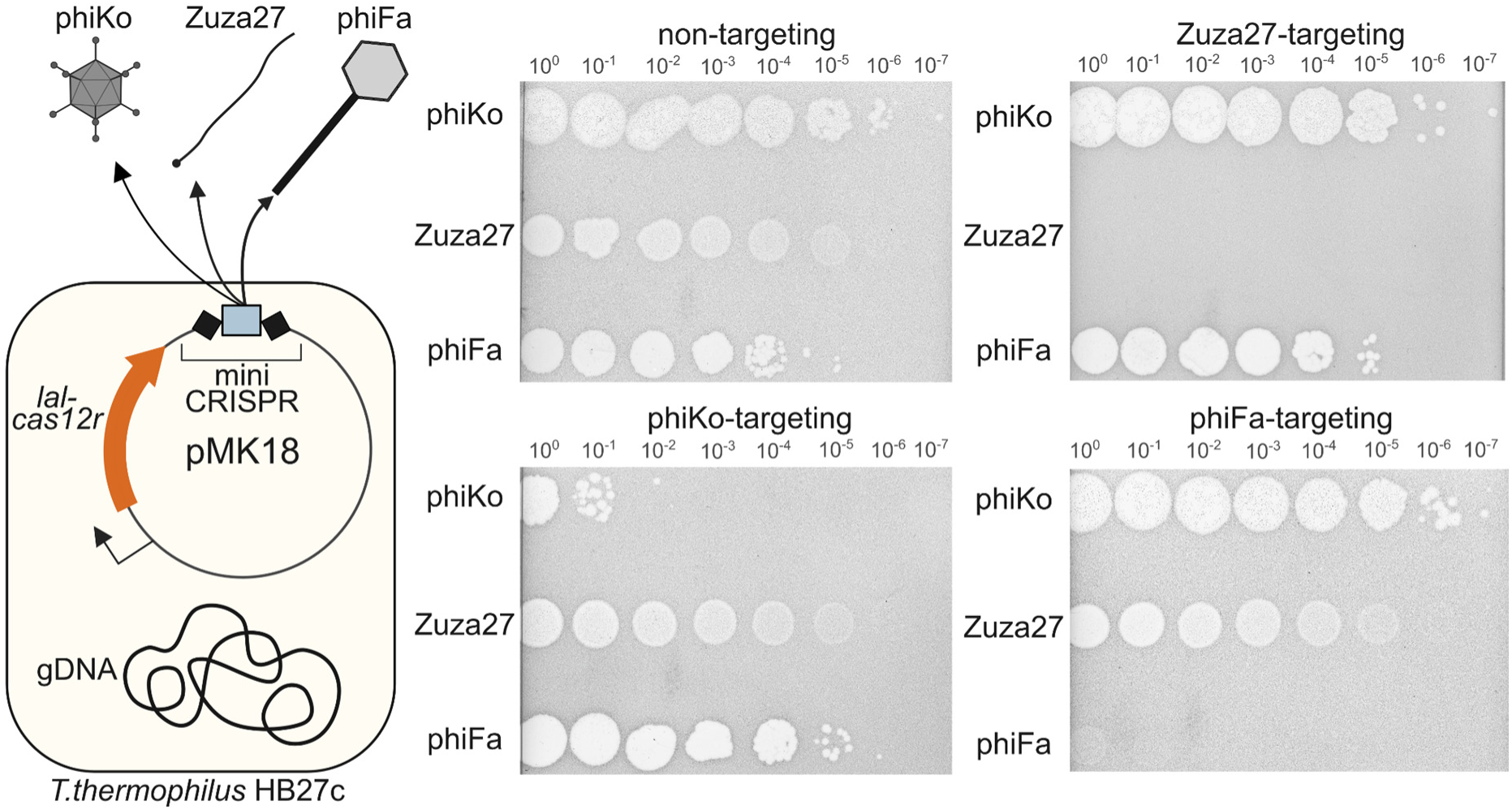
Lalka-derived LalCas12r–CRISPR system interferes with infection by diverse *T. thermophilus* phages. On the left, a schematic representation of the experimental setup is shown. 10-fold serial dilutions of phiKo, Zuza27 and phiFa phage lysates were spotted on the surface of lawns of *T. thermophilus* HB27c cells carrying the indicated phage-targeting or non-targeting control plasmids. Results of the overnight incubation are shown.

Despite robust interference observed in experiments with plasmid transformation and phage infection, rare individual *T. thermophilus* colonies transformed by GT or ST plasmids (escaper cells) and phage plaques on lawns of cells harboring PT plasmids (escaper phages) were observed. Escaper cells could have contained plasmids with inactivated Lalcas12r-CRISPR system or mutated target protospacer/PAM while escaper phages were expected to acquire mutations in target protospacer/PAM. To test whether plasmids in escaper cells accumulated mutations disabling LalCas12r interference, we extracted plasmids from several randomly selected escaper colonies and retransformed them into wild-type *T. thermophilus* HB27c cells. All escaper-derived plasmids transformed *T. thermophilus* HB27c cells with efficiency comparable to that of empty pMK18 vector purified from *T. thermophilus* HB27c cells (**Supplementary Fig. S10A**). We noticed that pMK18 purified from *T. thermophilus* HB27c re-transformed with a ∼100 times higher efficiency than the plasmid purified from *E. coli* cultures (**Supplementary Fig. S10A**). Although beyond the scope of the present study, we take this result as an indication that plasmids from *T. thermophilus* HB27c contain epigenetic modifications that protect them from *Thermus*-encoded restriction-modification systems. Indeed, *T. thermophilus* HB27c strain harbors bifunctional restriction endonuclease/methyltransferase Tth111II, which could be responsible for the observed effect (60).

Sequence analysis of plasmids purified from escaper cells revealed mutations either in mini-CRISPR arrays or in the *lalcas12r* gene. Some plasmids lacked spacers and contained only a single repeat unit, likely a result of recombination between repeats within the mini-CRISPR arrays. Mutations within the *lalcas12r* gene included small indels resulting in frameshifts, missense mutations, and an in-frame deletion that removed 28 amino acids from the N-terminal region of LalCas12r, as well as insertions of an *E. coli* IS4 element, which were likely acquired during plasmid propagation in *E. coli* cells (**Supplementary Fig. S10B**).

Individual phiKo phage escapers infected *T. thermophilus* HB27c cells harboring plasmids with phiKo-targeting spacers with the same efficiency as cells harboring the NT plasmid (**Supplementary Fig. S11A**). Amplification and sequencing of protospacer-containing genomic regions of phiKo escapers revealed single-nucleotide substitutions in the 5′-proximal segment of the protospacer (**Supplementary Fig. S11B**) indicating that this segment may represent a seed sequence (61). It should be noted that the targeted phiKo protospacer is located within the DNA polymerase gene, which must be essential for phage replication. This should thereby restrict the range of escaper mutations to those that do not compromise viral viability.

### Small RNAs associated with LalCas12r-CRISPR system

To identify RNAs utilized by LalCas12r-CRISPR system, a dual-spacer plasmid pMK18_*lalcas12r*–r_phiKo_r_phiFa_r encoding a *lalcas12r*-CRISPR locus with an array сontaining two spacers targeting phiKo and phiFa phages was constructed. The dual-spacer plasmid protected *T. thermophilus* HB27c cells against both phages, indicating that both spacers supply the effector with functional guide RNAs (**Supplementary Fig. S12A**). We used sRNA-Seq to identify small RNAs in *T. thermophilus* HB27c cells harboring dual-spacer plasmids containing either wild-type *lalCas12r* or catalytically inactive *dlalCas12r* genes. Additionally, small RNAs from the cells harboring empty pMK18 were sequenced. The sRNA-Seq reads were mapped to the *T. thermophilus* HB27c reference genome as well as to the pMK18_*lalcas12r*–r_phiKo_r_phiFa_r plasmid.

We observed two sRNA-Seq coverage peaks within the plasmid-borne CRISPR array **(Fig. 5A, Supplementary Fig. S12B)**. These peaks corresponded to 44 nt long RNA sequences consisting of a 26-nt repeat-derived 3′ segment and an 18-nt spacer-derived 5′ segment, originating either from the first or the second spacer. Additionally, we observed a coverage peak with a 26-nt repeat-derived 3′ segment followed by an 18 nt sequence downstream of the last repeat of the plasmid-borne CRISPR array. We therefore hypothesized that a single repeat may be sufficient for LalCas12r crRNA production. To test this hypothesis, we constructed a plasmid encoding the *lalcas12r* gene, the downstream intergenic sequence, and a single repeat followed by an 18 bp 5′ segment of the phiKo targeting spacer that protected cells against phiKo infection (above, **Fig. 4).** *T. thermophilus* HB27c cells harboring this plasmid were resistant to phiKo phage infection and sRNA-Seq analysis revealed a 44-nt crRNA comprising the last 26-nt of the repeat and the 18-nt downstream sequence **(Fig. 5B, Supplementary Fig. S12B**). Thus, a single repeat is indeed sufficient for generation of LalCas12r guide RNAs. Next, we assessed sRNA-Seq data of *T. thermophilus* HB27c cells infected by Lalka27a. Although we did not observe distinct sRNA-seq coverage peaks corresponding to the 5′ and 3′ ends of mature guide RNAs, we detected decreased sRNA-seq coverage depth at positions located 18 bp downstream of the repeat units (**Supplementary Fig. S12С**), which we take as evidence of crRNA production during the infection.

**Figure 5.**
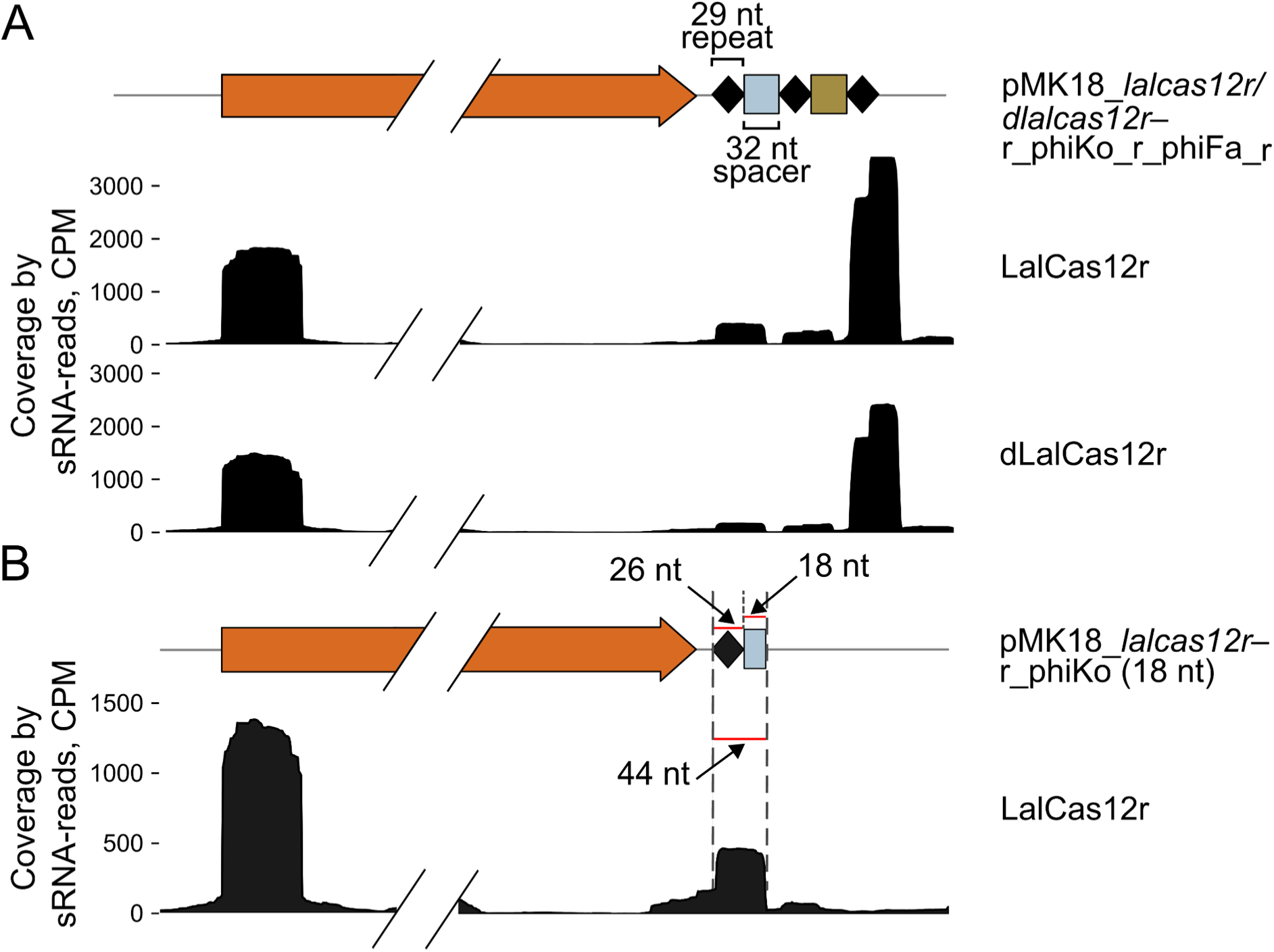
Guide RNAs in *T. thermophilus* HB27c cells carrying dual-spacer plasmids with *lalсas12r* or *dlalсas12r*. Coverage of the *lalcas12r/dlalcas12r*_mini-CRISPR locus by sRNA-Seq reads from RNA isolated from *T. thermophilus* HB27c cells harboring the pMK18_*lalcas12r/dlalcas12r*–r_phiKo_r_phiFa_r plasmid (**A**) or a plasmid encoding *lalcas12r* and a single repeat followed by the 18-nt spacer targeting phiKo phage (**B**). The LalCas12r variants (LalCas12r or dLalCas12r) and the type of the plasmid are shown to the right of the read coverage. For uncropped images see **Supplementary Fig. S12B**.

While the maturation of guide RNAs in some Type V CRISPR-Cas systems requires additional non-coding RNAs, i.e., trans-activating crRNA (tracrRNA) or short-complementarity untranslated RNA (scoutRNA) (62), the survey of sRNA-Seq reads of cells expressing *lalcas12r-*CRISPR loci did not reveal small RNAs beyond the crRNAs. Given that the tracrRNAs and scoutRNAs are known to be highly abundant (62, 63), the LalCas12r-CRISPR system apparently functions with crRNAs only.

In many Type V CRISPR-Cas systems and RNA-guided TnpB nucleases, the maturation of guide RNAs relies on the nuclease activity of the RuvC domain of Cas12/TnpB proteins (64–66). To test whether the RuvC domain of LalCas12r is involved in crRNA maturation, we sequenced small RNAs purified from *T. thermophilus* HB27c cells harboring dual-spacer plasmid encoding the dLalCas12r protein. While these cells were sensitive to the infection by both phages, they displayed the same set of small RNAs as cells harboring the plasmid encoding active LalCas12r (**Fig. 5A**, **Supplementary Fig. S12A**). Thus, the catalytic activity of the RuvC nuclease of LalCas12r is not required for guide maturation.

## DISCUSSION

In this work, we report the discovery of a novel Type V effector, Cas12r. We show that the plasmid-borne Lalka phage-derived Cas12r–CRISPR system interferes with plasmid transformation in *T. thermophilus* and protects these cells from infection by diverse phages when provided with appropriate spacers. Weblogo analysis and experimental data indicate that Cas12r recognizes and cleaves targets in a 5′ A-rich (ARG) PAM-dependent manner, in contrast to most other Cas12 nucleases, which preferentially recognize 5′ T-rich PAMs (63, 67–72). We determined that Cas12r crRNA is 44 nt long and consists of a 26-nt repeat-derived 3′ segment and an 18-nt spacer-derived 5′ segment. A single CRISPR repeat is sufficient for generation of LalCas12r guide RNAs, suggesting that crRNAs are generated by a ruler-like mechanism. Previously, small RNAs referred to as extraneous crRNAs, derived from marginal repeat units and flanking sequences outside the arrays, have been observed in various CRISPR–Cas systems. As these RNAs are incapable of mediating interference against invaders, they may hamper the immune response by sequestering Cas proteins. Accordingly, many CRISPR–Cas systems have independently evolved diverse mechanisms to suppress formation of extraneous crRNAs (73–75). The absence of such mechanisms in LalCas12r CRISPR systems likely reflects their recent evolutionary origin.

Although the immune response depends on the activity of Cas12r RuvC nuclease, this activity is dispensable for crRNA maturation. This is in contrast to several other Type V CRISPR–Cas systems and RNA-guided TnpB nucleases in which pre-crRNA processing occurs in the RuvC active site. The length of Cas12r (564 aa) falls within the range of the shortest known Type V effectors, such as Cas12f1 (400–700 aa) and Cas12n (506 aa) (76, 77). However in contrast to these proteins, Cas12r does not require tracrRNA for immune response. Considering the thermostability of Cas12r, its small size, lack of tracr/scout RNAs, and a rare PAM preference, it may become a useful tool for genome-editing in thermophilic organisms.

New Type V CRISPR–Cas variants typically emerge through association of transposon-encoded RNA-guided TnpB nucleases, the evolutionary ancestors of Cas12 effectors, with CRISPR arrays (32). LalCas12r and LlkCas12r evolved from a branch of TnpB nucleases for which association with CRISPR arrays has not been previously reported (**Fig. 6A**). A closer examination of this branch reveals that all TnpB proteins originated from thermophilic organisms (**Supplementary Table S8)**. We suggest that all Cas12r–CRISPR systems may also be thermophile-specific.

**Figure 6.**
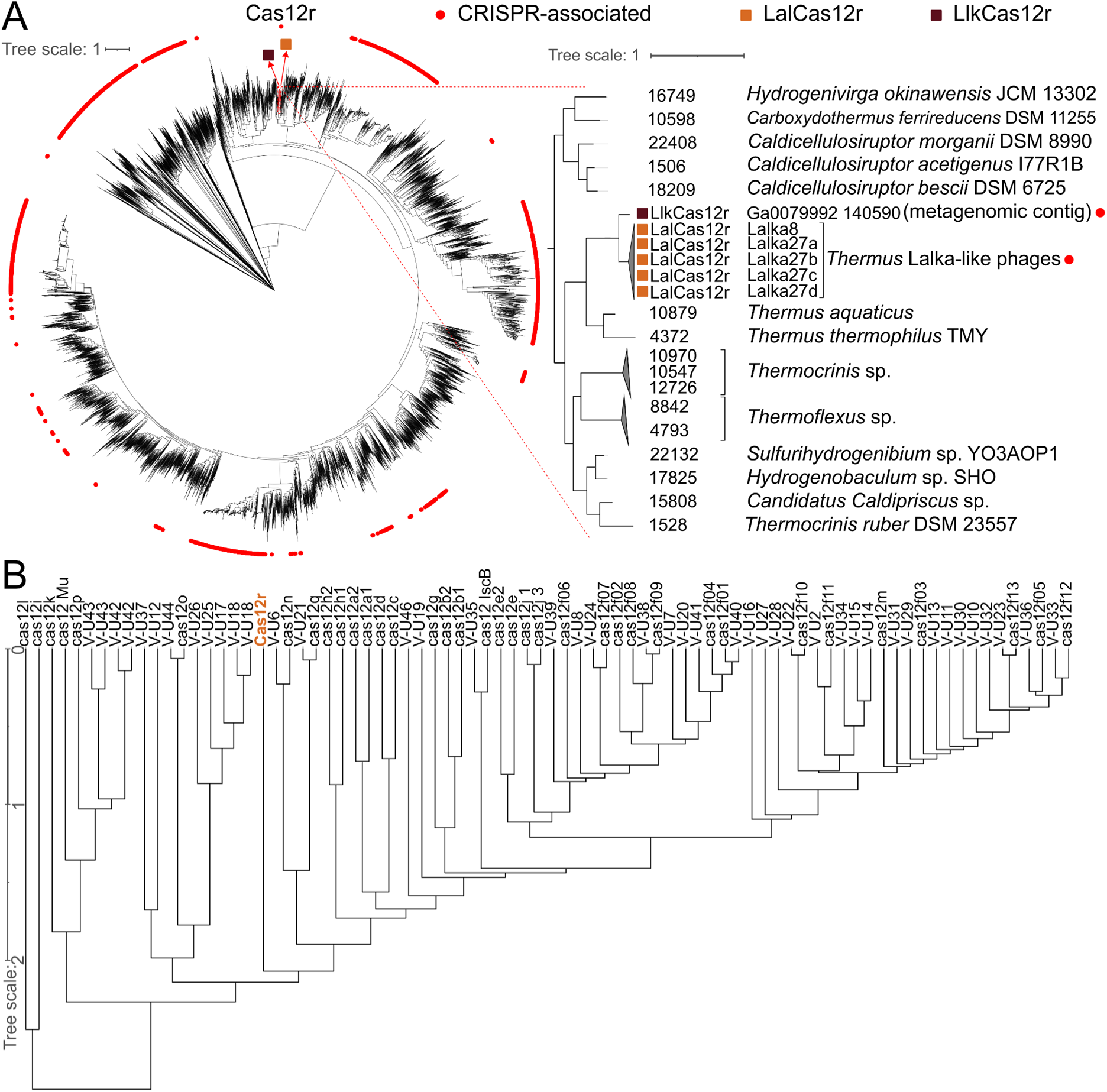
LalCas12r and LlkCas12r define a new subtype of Type V effectors. **(A)** A phylogenetic tree of TnpB proteins. Red circles on the outside indicate homologues associated with CRISPR arrays. The TnpB branch containing LalCas12r and LlkCas12r is colored red and is zoomed-into on the right. (**B)** A tree (UPGMA dendrogram based on profile-profile comparison scores) of the RuvC domains of Cas12r and known Cas12 subtypes.

To assess the evolutionary relationship between the Cas12r, known Cas12 effectors, we compared their RuvC-like nuclease domains. Previously established Cas12 RuvC profiles (78) were combined with the RuvC domain alignment of LalCas12r and LlkCas12r CRISPR-associated proteins (indicated as Cas12r on a dendrogram) **(Fig. 6B**). The relative distances within this dendrogram, comparable to or greater than those separating distinct type V subtypes, indicate that LalCas12r and LlkCas12r form an evolutionarily distinct cluster, supporting their classification as members of a novel subtype of type V effectors, as per established CRISPR-Cas classification criteria (78, 79).

The diversity of spacers within Lalka-encoded CRISPR arrays suggests an ongoing spacer acquisition process (**Supplementary Table S5**). The last spacer in Lalka27b, Lalka27c, and Lalka27d CRISPR arrays is identical, while the upstream spacers differ, suggesting that new spacers are inserted proximal to the *cas12r* gene. Since no adaptation genes or predicted ORFs with such functions are encoded in Lalka genomes, the adaptation process must rely on host functions. The type I-E CRISPR–Cas system is most widespread and present in ∼50% of *Thermus* genomes (**Supplementary Fig. S13)**. It relies on a 5′ AAG PAM (15), which overlaps with the PAM consensus sequence of the Cas12r-CRISPR system. Like the Cas12r–CRISPR repeat, the type I-E repeat is 29 bp long, and their sequences share common elements (**Supplementary Fig. S7**). Also, the average length of *Thermus* Type I-E spacers is 32 bp long (15) which is comparable to the Lalka-encoded 32-33 bp spacers. Thus, we tentatively propose that spacer acquisition by Cas12r–CRISPR might be mediated by the adaptation module of the *Thermus* type I-E CRISPR–Cas system.

Spacers from Lalka phage CRISPR arrays predominantly target integrative elements of *Thermus* that use the 3’ proximal elements of different host tRNA genes as attachment sites. Further, the integrases of *Thermus* IEs targeted by LalCas12r-CRISPR are very similar to those of Lalka phages (**Supplementary Fig. S8**). Finally, the Lalka-like prophage of *T. sp.* 93170 is inserted into the anticodon of the tRNA-Ala gene. Previously, the integrase encoded by ICETh2, a *Thermus* insertion element related to IEs targeted by Lalka phages, was shown to catalyze site-specific recombination between attachment sites in diverse tRNA genes (57). Given the similarity between the integrases encoded by Lalka phages, the IEs they target, and that of ICETh2, we propose that in the course of the infection, the Lalka integrase, a product of a middle gene, is capable of mobilizing the IE elements.

Targeting of integrated IEs by LalCas12r-CRISPR during the infection would be identical to host DNA targeting and does not appear to be an advantageous strategy for a phage capable of host lysogenization. Besides, if Lalka phage development required host targeting, one would expect the presence of spacers targeting host sequences other than IEs. It thus appears that Lalka phages specifically target the IEs of their hosts as part of inter-MGE competition that should involve IEs excised from the host genome. One possibility is that IEs may be mobilized during the infection and act as Lalka phage satellites. Phage satellites either lack or encode an incomplete set of structural genes, thus requiring a helper phage for the formation of infective virions. The genomes of phage satellites are much smaller than and distinct from the genomes of their helper phages. The genomes of IEs targeted by LalCas12r CRISPR are 6–15 kbs in length, comprising 8–21% of the length of Lalka phage genomes. The genomic organization of these elements, unlike those of known phage satellites, displays little conservation except for the presence of genes encoding an integrase and archaeo-eukaryotic primase-polymerase followed by a helicase gene. The ICETh2-like elements have the same organisation (80).

Phage genomes encode various defense and counterdefense systems (5, 81). In some closely related phages, different defense or counterdefence loci are encoded within identical genomic contexts (5, 82). For example, the genomes of some ICP1 phages encode a PLE-degrading nuclease in place of the Type I-F CRISPR-Cas locus (83). One of the discovered Cas12r-CRISPR loci, LlkCas12r-CRISPR, is apparently encoded by a Lalka-related bacteriophage (**Supplementary Fig. S6B**). Interestingly, while all Lalka Cas12r-CRISPR loci are located within the cluster of structural genes, the orientation of the LlkCas12r-CRISPR locus is opposite of that of the LalCas12r-CRISPR locus. This must be due to either by independent acquisition of LlkCas12r-CRISPR by a Lalka-related phage or by a rearrangement in the viral genome. Both scenarios imply that the cluster of structural genes in Lalka-related phages contain a recombination hot spot. Analysis of variable segments of Lalka-related phage genomes could reveal additional novel defense or counterdefense mechanisms in this location.

While LalCas12r CRISPR likely interferes with replication and/or propagation of targeted IEs, these elements may themselves interfere with phage development as they are enriched in diverse defense systems. These include a predicted *hicBA* toxin-antitoxin system in IE from *T. thermophilus* SNM1-7, a Type III restriction-modification system in IE from *Thermus sp.* H19.2, and a previously characterized (84) Class 1 OLD family nuclease from the IE element from *T. scotoductus* (CP001962.1:412381-431052). Class 1 OLD nucleases were shown to protect cells against phage infection (85, 86). Ongoing work in our laboratory on Lalka-like phages shall shed more light on their interactions with each other and the MGEs of their hosts.

## ACKNOWLEDGEMENTS

We would like to thank A. Rodionova for the work on the phiKo escapers during her internship. M.K. would like to thank K. Sayfulina for assistance during collection of environmental samples.

## SUPPLEMENTARY DATA FUNDING

A.T. and M.K. are supported by a grant from the Russian Science Foundation (25-24-00838) to M.K. A.T., M.K., and K.S. are also partially supported by a grant from the Russian Science Foundation (24-14-00181) to K.S.

## DATA AVAILABILITY

Raw sequencing data have been deposited with the National Center for Biotechnology Information Sequence Read Archive under BioProject ID PRJNA1452983.

