## Supplementary Figures and Tables for "A Unique Type V CRISPR-Cas System Encoded by a Group of *Thermus* Viruses": cas12r_manuscript_supplementary_figures_tables.pdf

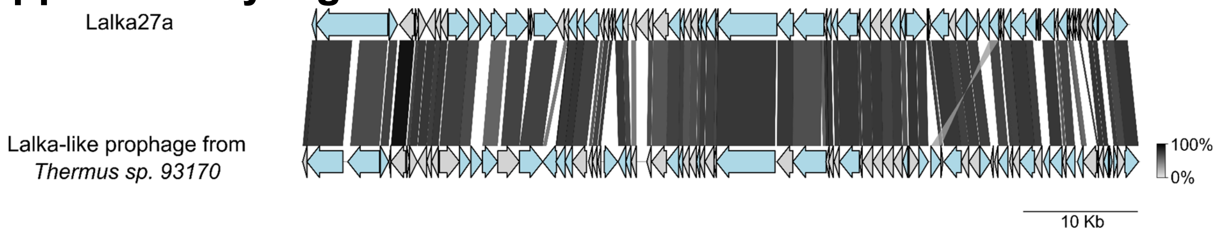

**Supplementary Figure S1.** Graphical alignment of the Lalka27a genome with the Lalka-like prophage region of *Thermus sp.* 93170 (NZ_JBPXNL010000001.1:1975845-2048918). Homologous gene or gene segments shared between Lalka27a and ORFs of the *Thermus sp.* 93170 Lalka-like prophage are connected by shading, with shading intensity corresponding to amino acid sequence identity, as shown in the bar at the bottom right.

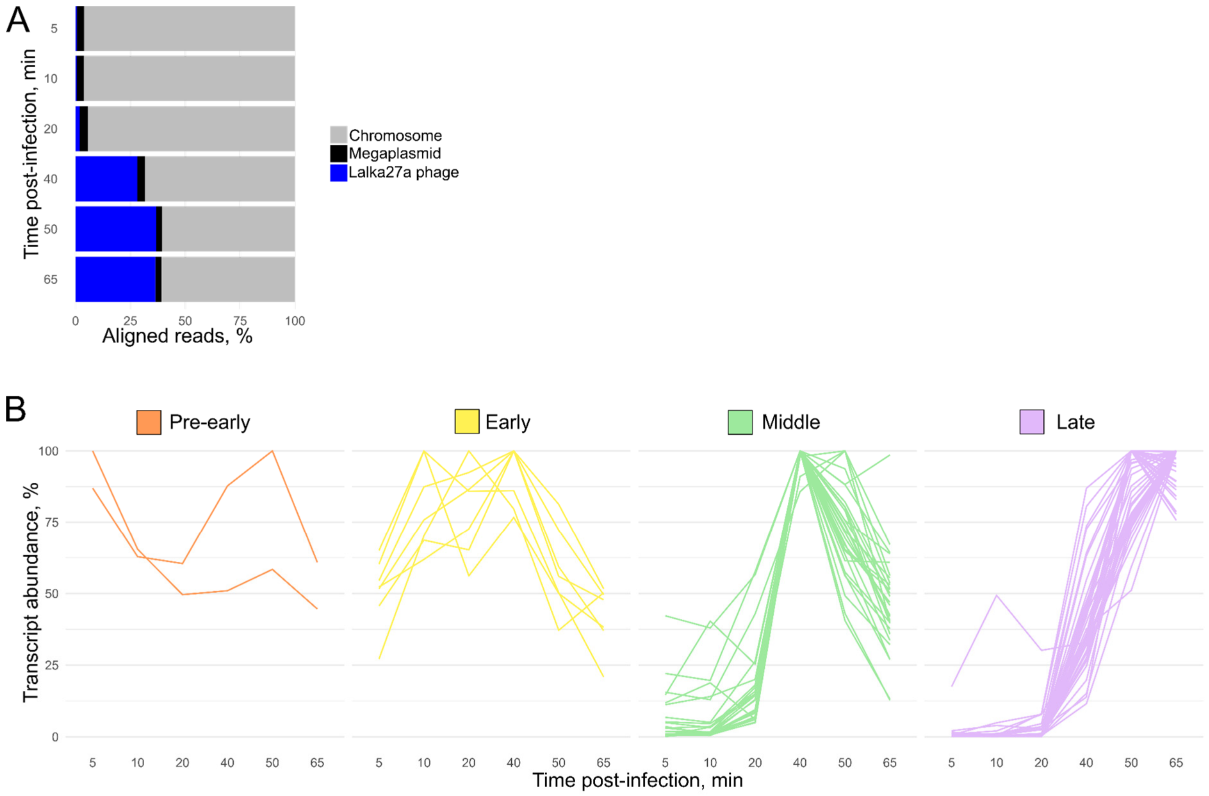

**Supplementary Figure S2.** Accumulation of Lalka27a transcripts throughout the infection.

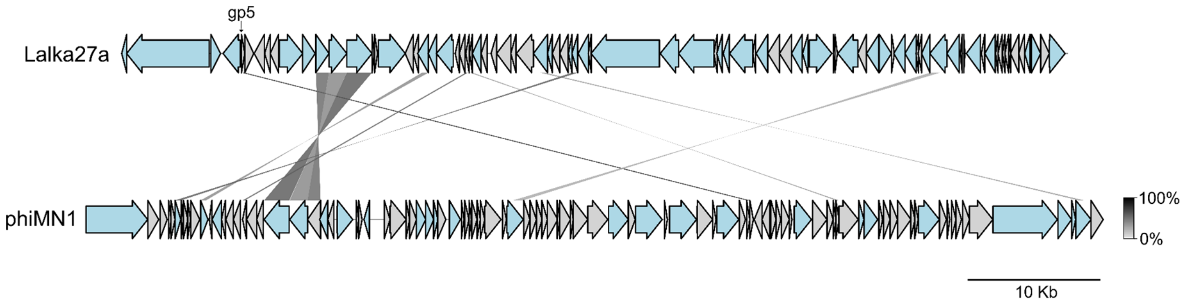

**Supplementary Figure S3.** Graphical alignment of the Lalka27a and phiMN1 phage genomes. For details see **Supplementary Fig. S1** legend.

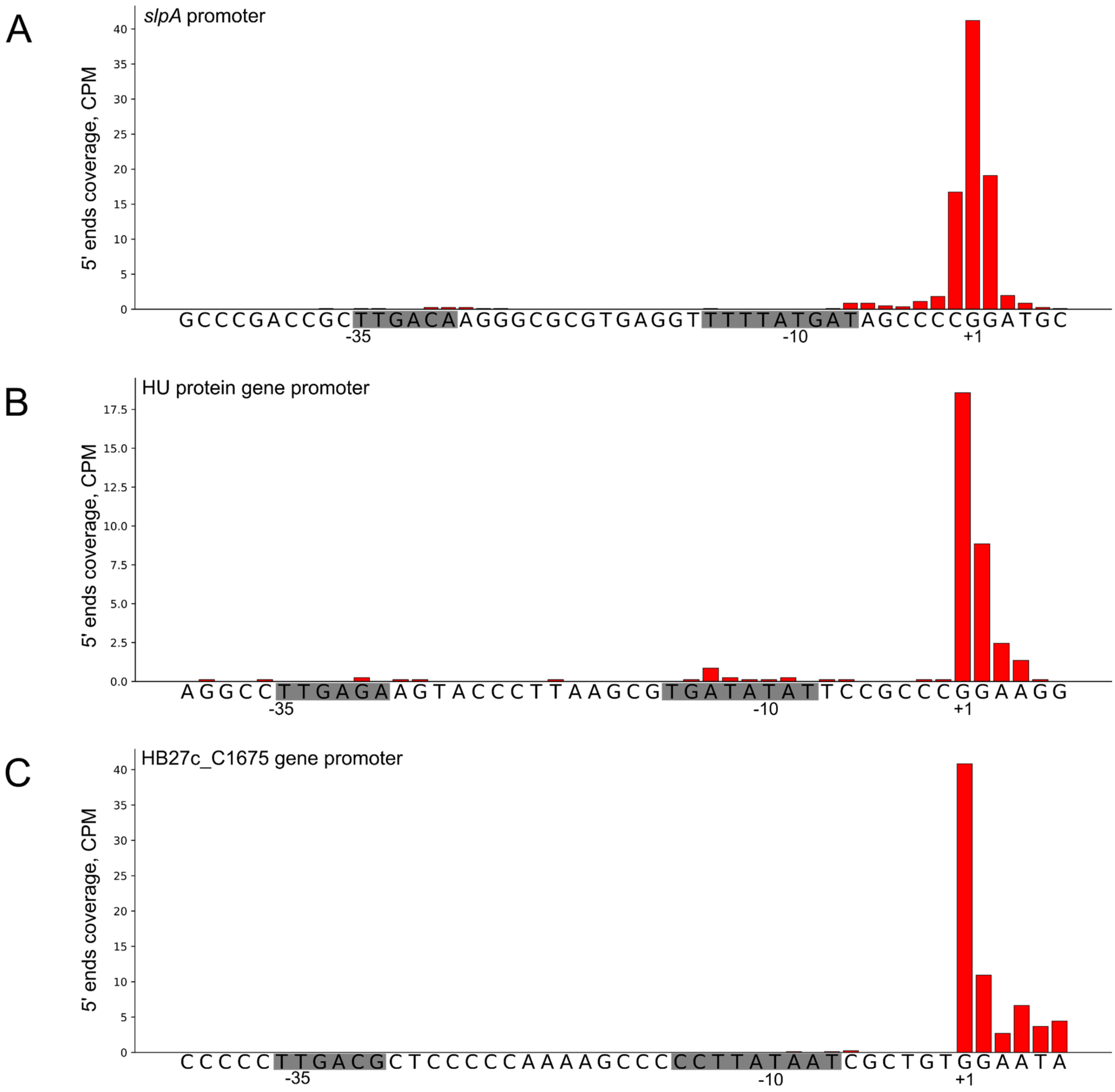

**Supplementary Figure S4.** CPM (counts per million) coverage of RNA 5' ends mapped onto promoter regions located upstream of the *Thermus slpA* gene (**A**), HU protein gene (**B**), and gene HB27c_C1675 encoding the ABC transporter substrate-binding protein (**C**). Promoter elements were predicted with the BPROM tool. sRNA-seq data obtained from *T. thermophilus* HB27c cells collected 40 min post-infection with Lalka27a were used.

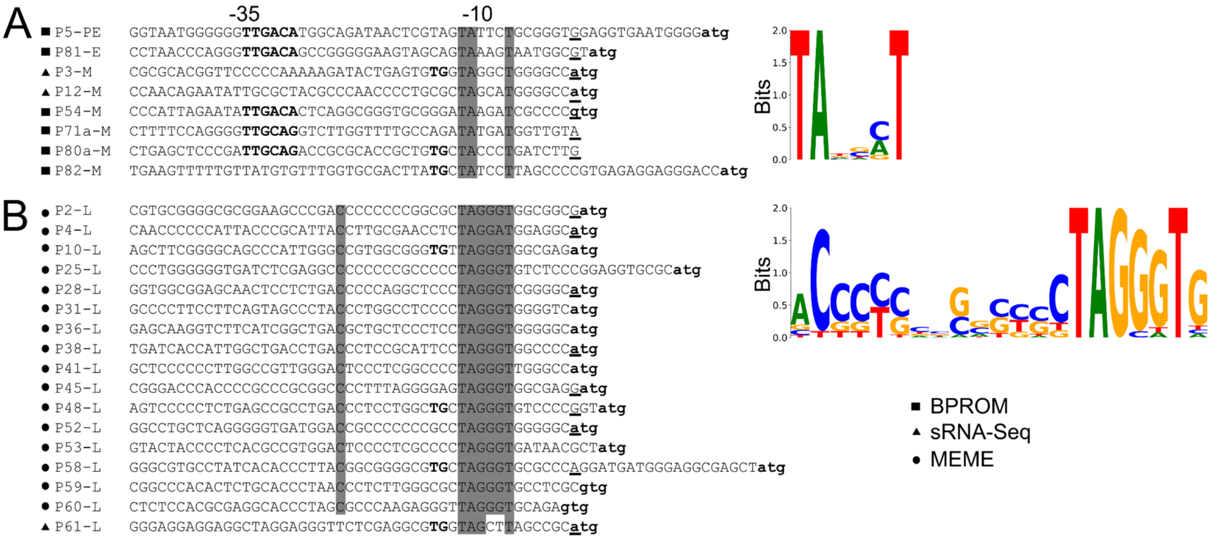

**Supplementary Figure S5. Predicted Lalka27a pre-early, early, and middle promoters** (**A**), and late promoters (**B**). Tools used for promoter prediction are indicated on the left by black squares (BPROM), triangles (sRNA-seq), and circles (MEME). Promoters identified by numbers corresponding to downstream Lalka27a genes are shown on the left. Promoter temporal classes are indicated: “PE” for pre-early, “E” for early, “M” for middle, and “L” for late promoters. Conserved nucleotide positions are highlighted in gray. The start codons of promoter-adjacent ORFs are shown in bold lowercase letters; the −35 promoter elements predicted with BPROM are shown in bold. Transcription start sites identified with sRNA-Seq are underlined. WebLogos of the conserved promoter regions are shown on the right.

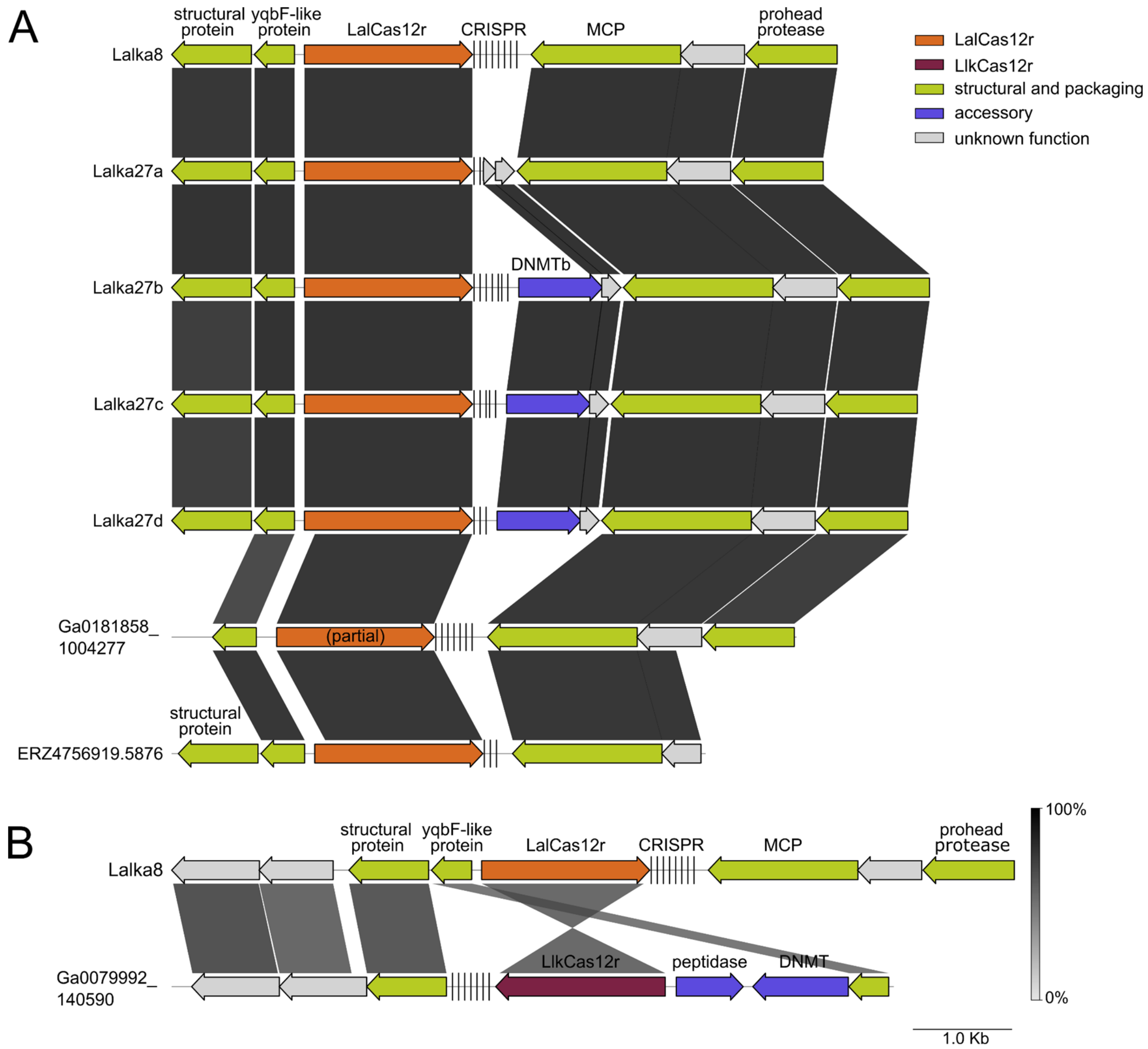

**Supplementary Figure S6. The diversity of Cas12r-CRISPR systems.** (**A, B**) Graphical alignment of genomic regions from Lalka phages and metagenomic contigs containing Cas12r-CRISPR systems. (**B**) Graphical alignment of LalCas12r and LlkCas12r loci and their genomic context. Homologous gene segments are connected by shading, with shading intensity corresponding to amino acid sequence identity, as shown in the bar at the bottom right.

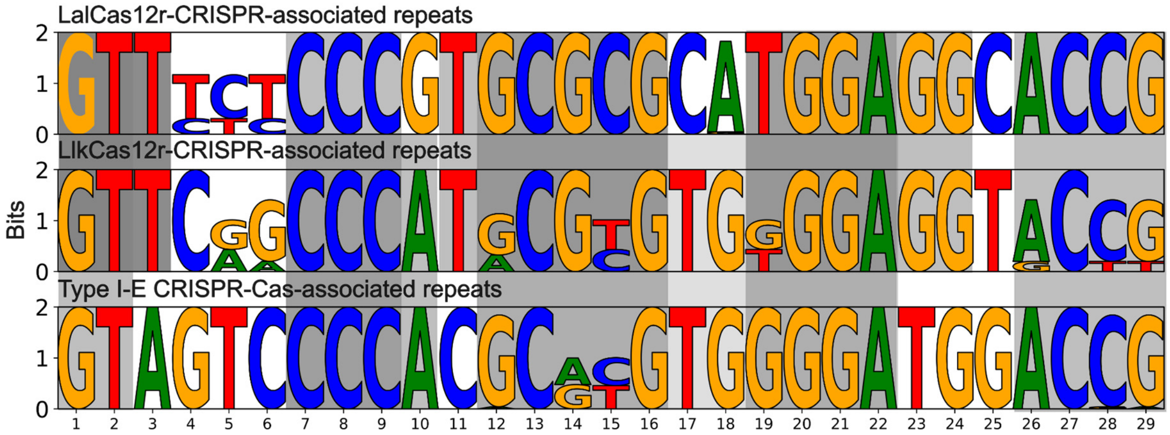

**Supplementary Figure S7.** Weblogos of repeat units associated with LalCas12r-CRISPR (top), LlkCas12r-CRISPR (middle) and *Thermus* Type I-E CRISPR-Cas (bottom) systems. Common elements shared between repeats are highlighted in grey. To build the weblogo for the Type I-E repeat units, the sequences of 153 Type I-E repeat units from *T. thermophilus* HB8, *T. brockianus* GE-1, and *Thermus sp.* 93170 were used.

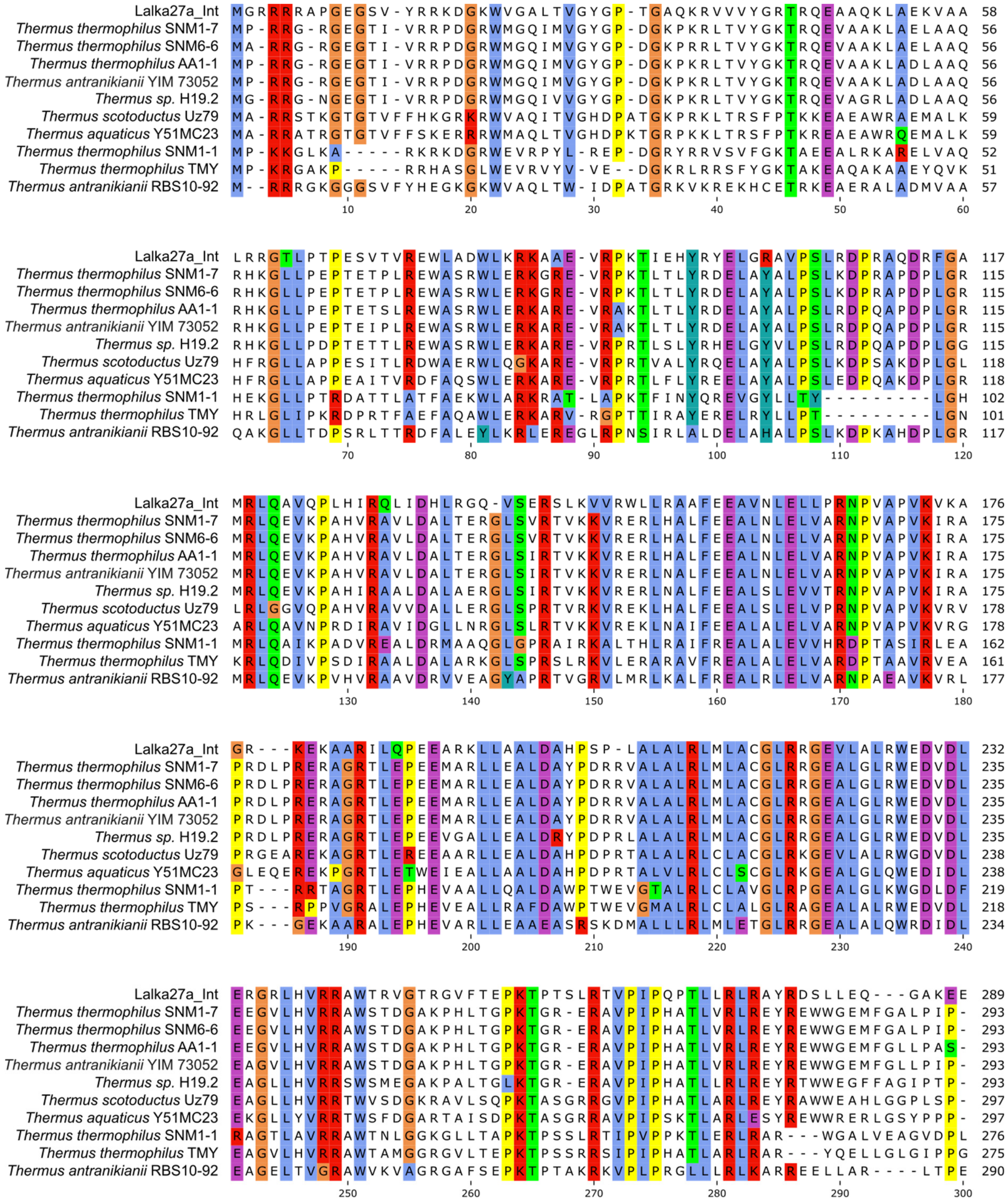
Lalka27a_Int MGRRRAPGEGSV-YRRKDGKWVGALTVGYGP-TGAQKRVVVYGRTRQEAAQKLAEKVAA 58
Thermus thermophilus SNM1-7 MP-RRG-RGEGTI-VRRPDGRWMGQIMVGYGP-DGKPKRLTVYGKTRQEVAAKLAELAAQ 56
Thermus thermophilus SNM6-6 MA-RRG-RGEGTI-VRRPDGRWMGQIMVGYGP-DGKPRRLTVYGKTRQEVAAKLAELAAQ 56
Thermus thermophilus AA1-1 MP-RRG-RGEGTI-VRRPDGRWMGQIMVGYGP-DGKPKRLTVYGKTRQEVAAKLADLAAQ 56
Thermus antranikianii YIM 73052 MP-RRG-RGEGTI-VRRPDGRWMGQIMVGYGP-DGKPKRLTVYGKTRQEVAAKLADLAAQ 56
Thermus sp. H19.2 MG-RRG-NGEGTI-VRRPDGRWMGQIVVGYGP-DGKPKRLTVYGKTRQEVAGRLADLAAQ 56
Thermus scotoductus Uz79 MA-RRSTKGTGTVFFHKGRKRWVAQITVGHDPATGKPRKLTRSFPTKKEAEAWRAEMALK 59
Thermus aquaticus Y51MC23 MA-RRATRGTGTVFFSKERRRWMAQLTVGHDPKTGRPKKLTRSFPTKREAEAWRQEMALK 59
Thermus thermophilus SNM1-1 MP-KKGLKA-----RKRKDGRWEVRPYL-REP-DGRYRRVSVFGKTAEEALRKARELVAQ 52
Thermus thermophilus TMY MP-KRGAKP-----RRHASGLWEVRVYV-VE--DGKRLRRSFYGKTAKEAQAKAAEYQVK 51
Thermus antranikianii RBS10-92 M--RRRGKGGGSVFYHEGKGKWVAQLTW-IDPATGRKVKREKHCETRKEAERALADMVAA 57
10 20 30 40 50 60
Lalka27a_Int LRRGTLPTPESVTVREWLADWLKRKAAE-VRPKTIEHYRYELGRAVPSLRDPRAQDRFGA 117
Thermus thermophilus SNM1-7 RHKGLLPEPTETPLREWASRWLERKGRE-VRPKTLTLYRDELAYALPSLKDPRAPDPLGR 115
Thermus thermophilus SNM6-6 RHKGLLPEPTETPLREWASRWLERKGRE-VRPKTLTLYRDELAYALPSLKDPRAPDPLGR 115
Thermus thermophilus AA1-1 RHKGLLPEPTETSLREWASRWLERKARE-VRAKTLTLYRDELAYALPSLRDPQAPDPLGR 115
Thermus antranikianii YIM 73052 RHKGLLPEPTEIPLREWASRWLERKARE-VRAKTLTLYRDELAYALPSLRDPQAPDPLGR 115
Thermus sp. H19.2 RHKGLLPDPTETTLREWASRWLERKARE-VRPRTLSLYRHELGYVLPSLRDPQAPDPLGG 115
Thermus scotoductus Uz79 HFRGLLAPPESITLRDWAERWLQGKARE-VRPRTVALYRQELAYALPSLKDPSAKDPLGL 118
Thermus aquaticus Y51MC23 HFRGLLAPPEAITVRDFAQSWLERKARE-VRPRTLFLYREELAYALPSLEDPQAKDPLGR 118
Thermus thermophilus SNM1-1 HEKGLLPTRDATTLATFAEKWLARKRAT-LAPKTFINYQREVGYLLTY---------LGH 102
Thermus thermophilus TMY HRLGLIPKRDPRTFAEFAQAWLERKARV-RGPTTIRAYERELRYLLPT---------LGN 101
Thermus antranikianii RBS10-92 QAKGLLTDPSRLTTRDFALEYLKRLEREGLRPNSIRLALDELAHALPSLKDPKAHDPLGR 117
70 80 90 100 110 120
Lalka27a_Int MRLQAVQPLHIRQLIDHLRGQ-VSERSLKVVRWLLRAAFEEAVNLELLPRNPVAPVKVKA 176
Thermus thermophilus SNM1-7 MRLQEVKPAHVRAVLDALTERGLSVRTVKKVRERLHALFEEALNLELVARNPVAPVKIRA 175
Thermus thermophilus SNM6-6 MRLQEVKPAHVRAVLDALTERGLSVRTVKKVRERLHALFEEALNLELVARNPVAPVKIRA 175
Thermus thermophilus AA1-1 MRLQEVKPAHVRAVLDALTERGLSIRTVKKVRERLHALFEEALNLELVARNPVAPVKIRA 175
Thermus antranikianii YIM 73052 MRLQEVKPAHVRAVLDALTERGLSIRTVKKVRERLNALFEEALNLELVARNPVAPVKIRA 175
Thermus sp. H19.2 MRLQEVKPAHIRAALDALAERGLSIRTVKKVRERLNALFEEALSLEVVTRNPVAPVKIRA 175
Thermus scotoductus Uz79 LRLGGVQPAHVRAVVDALLERGLSPRTVRKVREKLHALFEEALSLELVPRNPVAPVKVRV 178
Thermus aquaticus Y51MC23 ARLQAVNPRDIRAVIDGLLNRGLSLRTVKKVREKLNAIFEEALALELVARNPVAPVKVRG 178
Thermus thermophilus SNM1-1 MRLQAIKPADVREALDRMAAQGLGPRAIRKALTHLRAIFREALALEVVHRDPTASIRLEA 162
Thermus thermophilus TMY KRLQDIVPSDIRAALDALARKGLSPRSLRKVLERARAVFREALALELVARDPTAAVRVEA 161
Thermus antranikianii RBS10-92 MRLQEVKPVHVRAAVDRVVEAGYAPRTVGRVLMRLKALFREALRLELVARNPAEAVKVRL 177
130 140 150 160 170 180
Lalka27a_Int GR---KEKAARILQPEEARKLLAALDAHPSP-LALALRLMLACGLRRGEVLALRWEDVDL 232
Thermus thermophilus SNM1-7 PRDLPRERAGRTLEPEEMARLLEALDAYPDRRVALALRLMLACGLRRGEALGLRWEDVDL 235
Thermus thermophilus SNM6-6 PRDLPRERAGRTLEPEEMARLLEALDAYPDRRVALALRLMLACGLRRGEALGLRWEDVDL 235
Thermus thermophilus AA1-1 PRDLPRERAGRTLEPEEMARLLEALDAYPDRRVALALRLMLACGLRRGEALGLRWEDVDL 235
Thermus antranikianii YIM 73052 PRDLPRERAGRTLEPEEMARLLEALDAYPDRRVALALRLMLACGLRRGEALGLRWEDVDL 235
Thermus sp. H19.2 PRDLPRERAGRTLEPEEVGALLEALDRYPDPRLALALRLMLACGLRRGEALGLRWEDVDL 235
Thermus scotoductus Uz79 PRGEAREKAGRTLEREEAARLLEALDAHPDPRTALALRLCLACGLRKGEVLALRWGDVDL 238
Thermus aquaticus Y51MC23 GLEQEREKPGRTLETWEIEALLAALDAHPDPRTALVLRLCLSCGLRKGEALGLQWEDIDL 238
Thermus thermophilus SNM1-1 PT---RRTAGRTLEPHEVAALLQALDAWPTWEVGTALRLCLAVGLRPGEALGLKWGDLDF 219
Thermus thermophilus TMY PS---RPPVGRALEPHEVEALLRAFDAWPTWEVGMALRLCLALGLRAGEALALRWEDVDL 218
Thermus antranikianii RBS10-92 PK---GEKAARALEPHEVARLLEAAEASRSKDMALLLRLMLETGLRRGEALALQWRDIDL 234
190 200 210 220 230 240
Lalka27a_Int ERGRLHVRRAWTRVGTRGVFTEPKTPTSLRTVPIPQPTLLRLRAYRDSLLEQ---GAKEE 289
Thermus thermophilus SNM1-7 EEGVLHVRRAWSTDGAKPHLTGPKTGR-ERAVPIPHATLVRLREYREWWGEMFGALPIP- 293
Thermus thermophilus SNM6-6 EEGVLHVRRAWSTDGAKPHLTGPKTGR-ERAVPIPHATLVRLREYREWWGEMFGALPIP- 293
Thermus thermophilus AA1-1 EEGVLHVRRAWSTDGAKPHLTGPKTGR-ERAVPIPHATLVRLREYREWWGEMFGLLPAS- 293
Thermus antranikianii YIM 73052 EAGVLHVRRAWSTDGAKPHLTGPKTGR-ERAVPIPHATLVRLREYREWWGEMFGLLPIP- 293
Thermus sp. H19.2 EAGLLHVRRSWSMEGAKPALTGLKTGR-ERAVPIPHATLLRLREYRTWWEGFFAGIPTP- 293
Thermus scotoductus Uz79 EAGLLHVRRTWVSDGKRAVLSQPKTASGRRGVPIPHATLARLREYRAWWEAHLGGPLSP- 297
Thermus aquaticus Y51MC23 EKGLLYVRRTWSFDGARTAISDPKTASGRRAVPIPSKTLARLESYREWWRERLGSYPPP- 297
Thermus thermophilus SNM1-1 RAGTLAVRRAWTNLGGKGLLTAPKTPSSLRTIPVPPKTLERLRAR---WGALVEAGVDPL 276
Thermus thermophilus TMY EAGVLHVRRAWTAMGGRGVLTEPKTPSSRRSIPIPHATLARLRAR---YQELLGLGIPPG 275
Thermus antranikianii RBS10-92 EAGELTVGRAWVKVAGRGAFSEPKTPTAKRKVPLPRGLLLRLKARREELLAR----LTPE 290
250 260 270 280 290 300

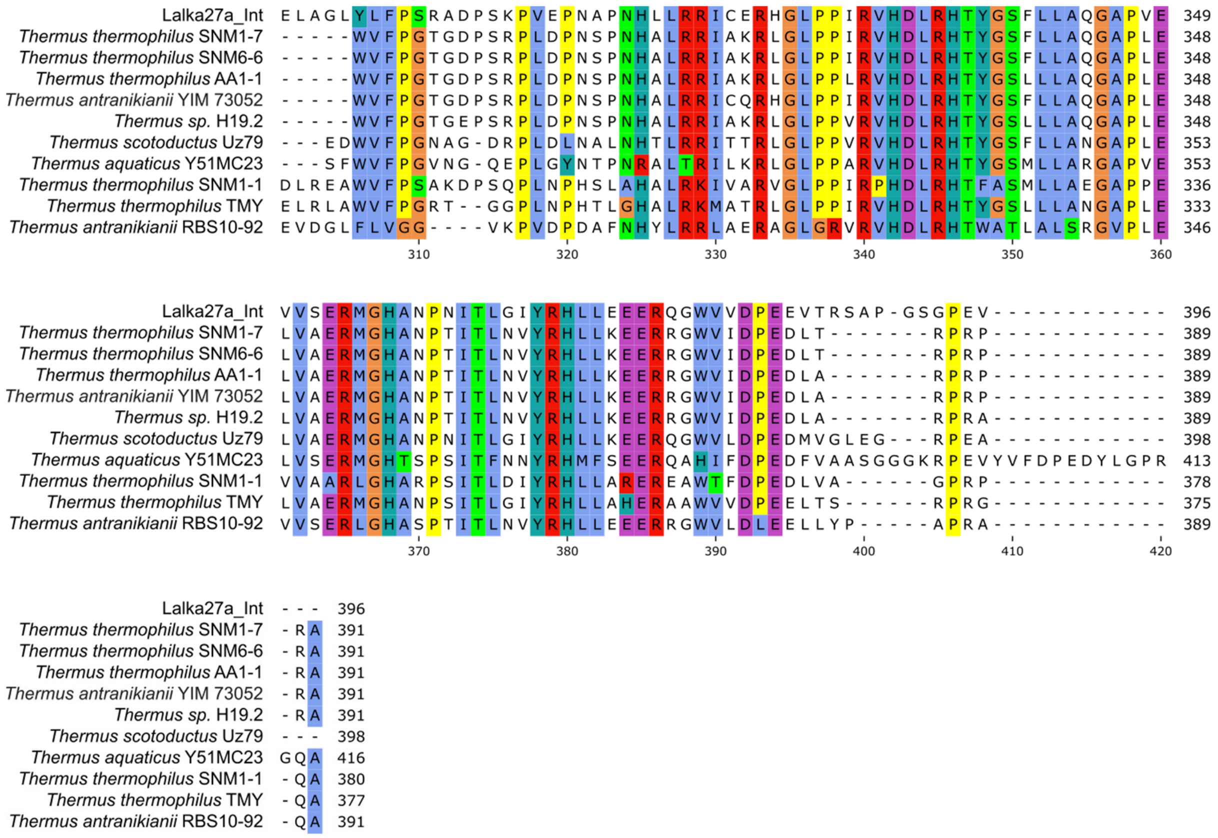

**Supplementary Figure S8.** Multiple alignment of integrase amino acid sequences from Lalka27a phage and LalCas12r–CRISPR-targeted IEs.

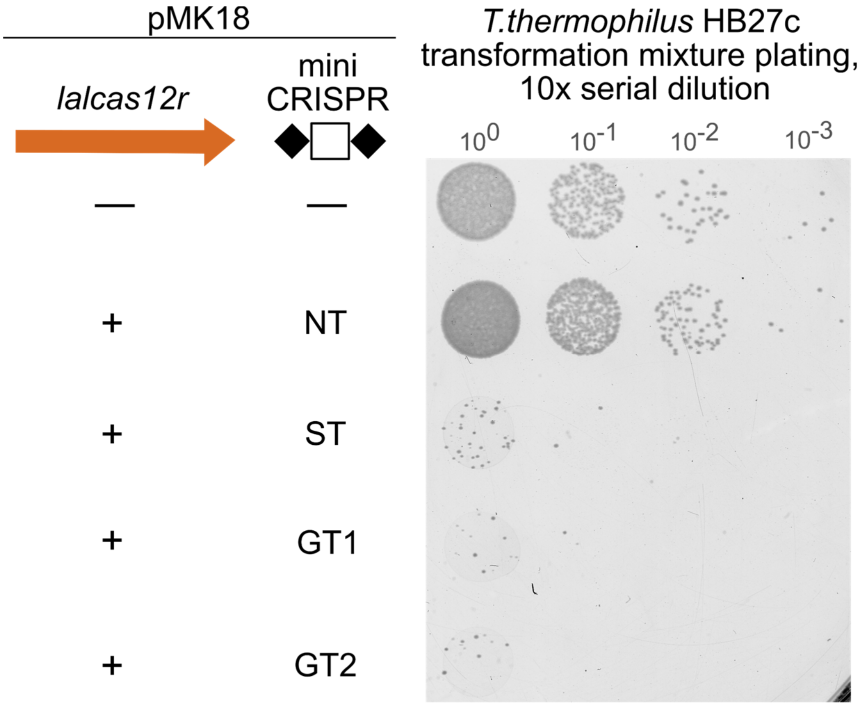

**Supplementary Figure S9. Lalka-derived Cas12r–CRISPR system interferes with plasmid transformation in *T. thermophilus* HB27c.** Overnight growth of 10-fold serial dilutions of *T. thermophilus* HB27c cells transformed with pMK18 or pMK18-derived plasmids encoding *lalcas12r* and a mini-CRISPR array carrying genome-targeting (GT1, GT2), self-targeting (ST), or non-targeting (NT) spacers on selective medium. Spacers from ST, GT1, and GT2 plasmids matched protospacers, which contained, at their 5′ ends, distinct 8-nt sequences that included a putative ARG PAM.

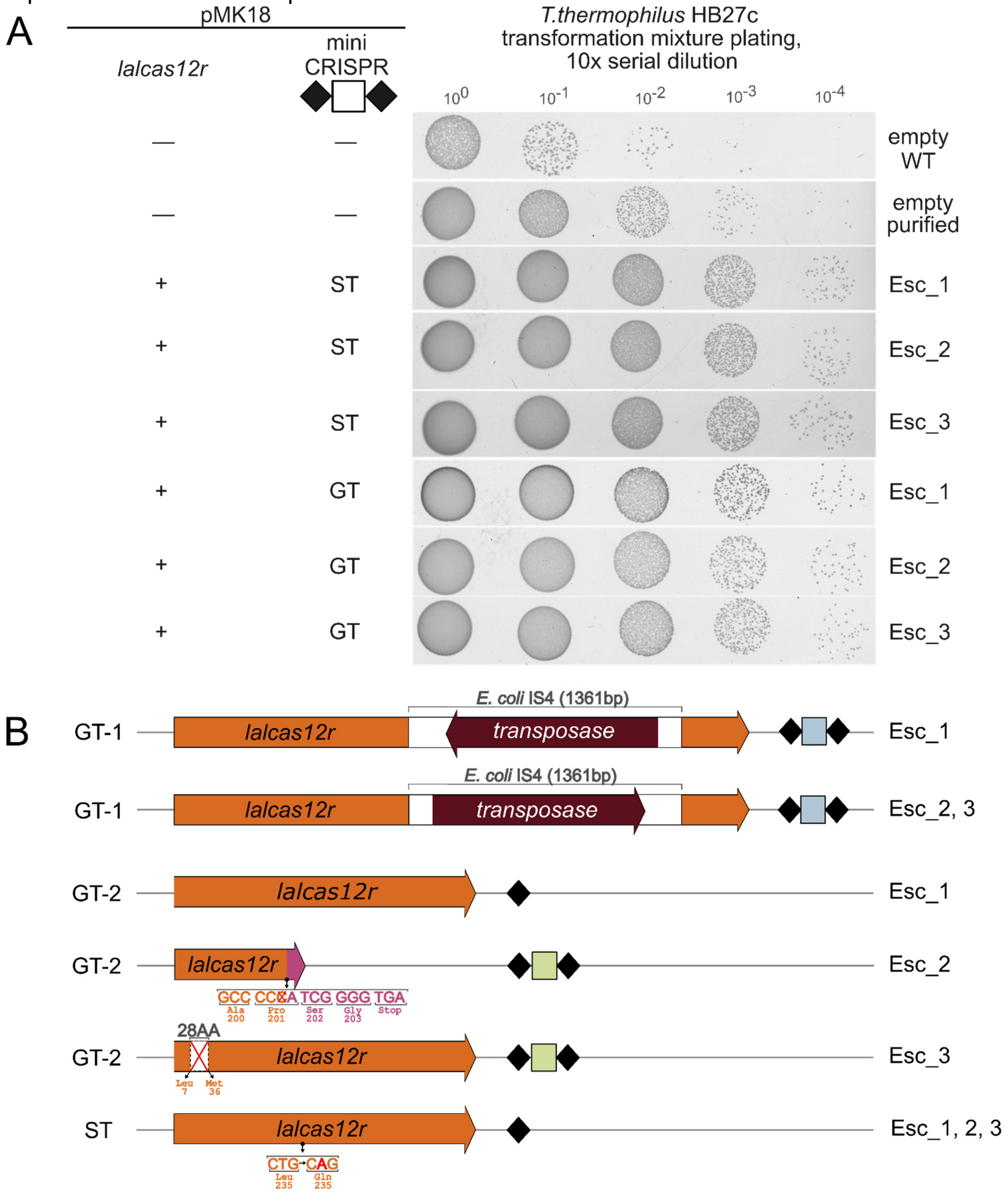

**Supplementary Figure S10. Escaper cells contain plasmids with mutations in mini-CRISPR arrays and/or in the *lalcas12r* gene.** (**A**) The overnight growth of aliquots of 10-fold serial dilutions of *T. thermophilus* HB27c transformation mixtures on selective medium. The GT and ST plasmids purified from three independent escaper cells colonies (Esc_1-3) were retransformed into *T. thermophilus* HB27c stock cultures; "empty WT" refers to pMK18 vector purified from *E. coli*, whereas "empty purified" refers to pMK18 purified from *T. thermophilus* HB27c. (**B**) Schematic representation of results of sequence analysis of GT and ST escaper plasmids. GT-1 plasmid escaper encodes the *lalcas12r* gene with the *E. coli* IS4 element inserted in different orientations. One GT-2 plasmid escaper (Esc_1) lacked the spacer and contained only one repeat unit. The second GT-2 escaper (Esc_2) carried a single-nucleotide deletion in the Pro201 codon (CCC → CCA), resulting in a frameshift and almost immediate truncation of LalCas12r due to formation of a premature stop codon after Gly203. The third GT-2 escaper (Esc_3) carried an in-frame deletion leading to the removal of 28 amino acids in the N-terminal segment of LalCas12r (between Leu7 and Met36). Mini-CRISPR arrays of all ST plasmid escapers lacked the spacer and contained only a single repeat unit. In addition, all ST plasmid escapers encoded the *lalcas12r* gene with a nonsynonymous mutation resulting in a Leu235→Gln substitution (CTG → CAG).

A

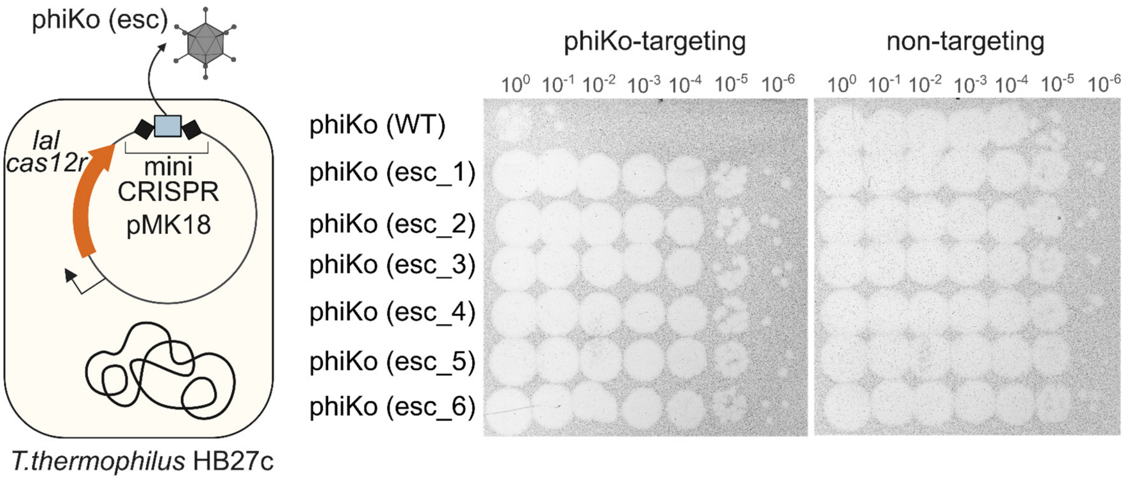

B

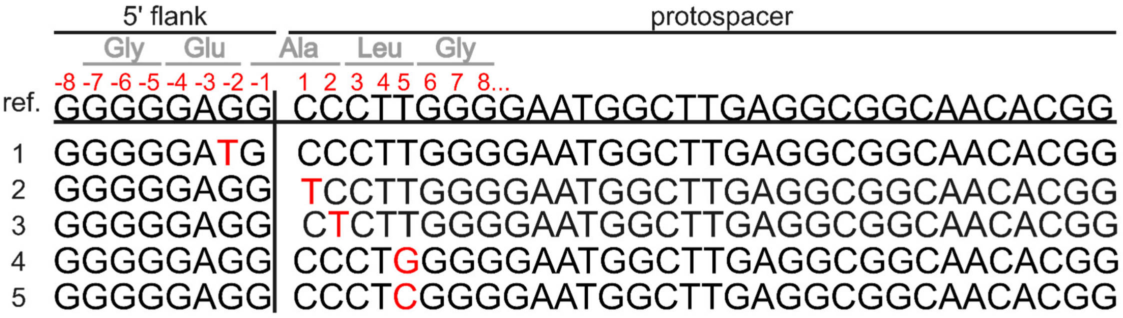

C

| Line | Mutation | Effect |
|---|---|---|
| 1 | G-2 ->T | Glu -> Asp |
| 2 | C1 ->T | Ala -> Val |
| 3 | C2 ->T | Synonymous mutation |
| 4 | T5 ->G | Synonymous mutation |
| 5 | T5 ->C | Synonymous mutation |

**Supplementary Figure S11. Escaper phages harbour mutations in target protospacer/PAM.** (**A**) On the left, a schematic representation of the experimental setup is shown. 10-fold serial dilutions of phiKo phage escaper lysates were spotted on the surface of lawns of *T. thermophilus* HB27c cells carrying the phiKo-targeting or NT plasmids. Results of overnight growth are shown. Spacers encoded in the mini-CRISPR arrays are indicated above the dilution series. Lysate types are shown to the left of the dilution series. "WT" denotes the stock phiKo phage lysate, "Esc" denotes lysates derived from individual phiKo phage escaper plaques. Numbers indicate experimental replicates. (**B**) Sequencing results of protospacer-containing genomic regions of phiKo phage escapers. "ref." corresponds to the reference phiKo phage sequence. Lines 1–5 correspond to escaper sequences. Nucleotide substitutions are shown in red. The sequence in line 1 contains a single nucleotide substitution at position −2 in the region adjacent to the 5′ end of the protospacer (putative PAM region). Sequences in lines 2–5 contain single nucleotide substitutions within the putative seed regions of the protospacers. (**C**) The table shows the effects caused by escaper mutations on the protein sequence.

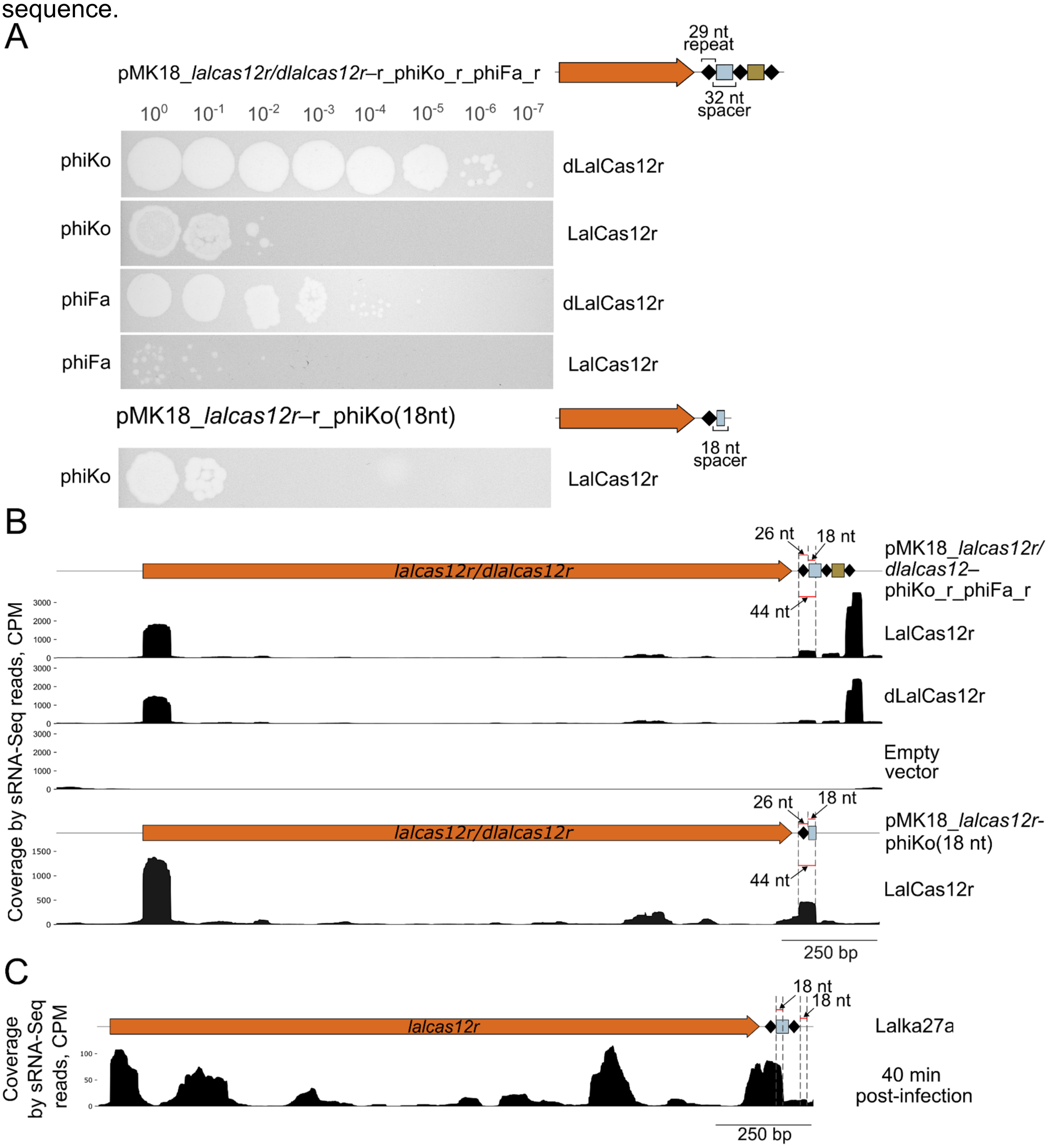

**Supplementary Figure S12. Maturation of three distinct guide RNAs in *T. thermophilus* HB27c cells carrying the dual-spacer plasmid with *lalcas12r* or *dlalcas12r*.** (**A**) 10-fold serial dilutions of phiKo/phiFa phage lysates were spotted on the surface of lawns of *T. thermophilus* HB27c cells carrying a dual-spacer plasmid with *lalcas12r/dlalcas12r* or a plasmid encoding *lalcas12r* and a single repeat followed by the 18-nt sequence targeting the phiKo phage. Results of overnight growth are shown. The type of the plasmid is shown above the dilution series. The lysate type is shown to the left of the dilution series. The LalCas12r variant is shown to the right of the dilution series. (**B**) Coverage of the *lalcas12r*_mini-CRISPR locus by sRNA-Seq reads from *T. thermophilus* HB27c cells harboring pMK18_*lalcas12r/dlalcas12r*–r_phiKo_r_phiFa_r plasmids, an empty pMK18 vector, or a plasmid encoding *lalcas12r* and a single repeat followed an 18-nt phiKo targeting sequence.

LalCas12r variants (LalCas12r or dLalCas12r) and the type of the plasmid are shown to the right of the read coverage. (**C**) Coverage of the Lalka27a-encoded *cas12r*_CRISPR locus by reads from sRNA-Seq of *T. thermophilus* HB27c cells infected with Lalka27a phage. sRNA-seq data obtained from *T. thermophilus* HB27c cells collected 40 min post-infection with Lalka27a are shown.

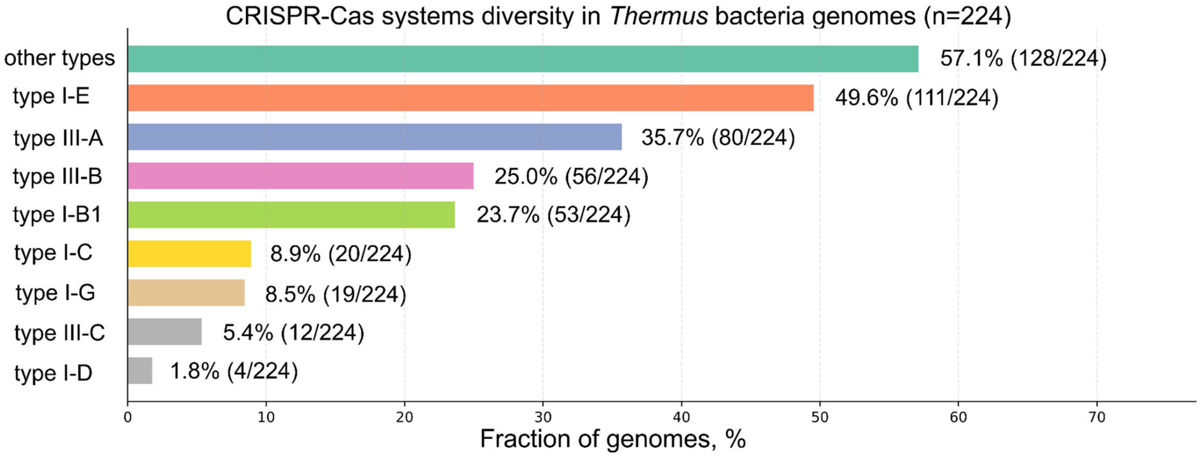

**Supplementary Figure S13.** CRISPR–Cas systems diversity in *Thermus* bacteria.

### Supplementary tables

**Supplementary Table S1.** Oligonucleotides used in the study.

| Name | Sequence 5′ to 3′ |
|---|---|
| Construction of plasmids encoding the LalCas12r and a mini-CRISPR array | |
| cas12r_putprom_F | TTATATGGTCTCGGGAGCACCTGCTTGTCCGAGTCGACTTGACACTCAGGCGGGTGCGGGATAAGATCGCC |
| cas12r_R | ATATAAGGTCTCAAGCGCACCTGCGGCGGCACGGGAGAAACACATGACCAGCACGCTCAAAACACCTCC |
| lacZ_F | TTATATGGTCTCGGGAGCACCTGCTTGTGTGCGCGCATGGAGGCACCGAGAGACGGCAGCTGGCACGACAGG |
| lacZ_R | ATATAAGGTCTCAAGCGCACCTGCGGCGAAACAGAGACGTTTGTCACAGCTTGTCTGTAAGCGG |
| repeat_F | GGAGATACACCTGCTTGTGTTTCTCCCGTGCGCGCATGGAGGCACCGGTCGACGCCCCTTCGCAGGTGTTAT |
| repeat_R | AGCGATAACACCTGCGAAGGGGCGTCGACCGGTGCCTCCATGCGCGCACGGGAG |

| | |
|---|---|
| | AAACACAAGCAGGTGTAT |
| Oligonucleotide sequences used for the construction of plasmids carrying mini-CRISPR arrays with different spacers | |
| GT1_spacer_F | ACCGAGCTCGTGGTCCGGGCCGCCCTC AAGCCCATC |
| GT1_spacer_R | AAACGATGGGCTTGAGGGCGGCCCGGA CCACGAGCT |
| GT2_spacer_F | TTCTCCTTGTCCTCGGCCTCTTGGTCAT GGTC |
| GT2_spacer_R | GACCATGACCAAGAGGCCGAGGACAAG GAGAA |
| ST_spacer_F | ACCGGAAGATCTGATTGCTTAACTGCTT CAGTTAAG |
| ST_spacer_R | AAACCTTAACTGAAGCAGTTAAGCAATC AGATCTTC |
| NT_spacer_F | ACCGCATTTCAGTCAGTTGCTCAATGTA CCTATAAC |
| NT_spacer_R | AAACGTTATAGGTACATTGAGCAACTGA CTGAAATG |
| phiKo-targeting_spacer_F | ACCGCCCTTGGGGAATGGCTTGAGGCG GCAACACGG |
| phiKo-targeting_spacer_R | AAACCCGTGTTGCCGCCTCAAGCCATTC CCCAAGGG |
| phiFa-targeting_spacer_F | ACCGGTGAACCCGTCCAGACCATCGGC GAATACGAA |
| phiFa-targeting_spacer_R | AAACTTCGTATTCGCCGATGGTCTGGAC GGGTTCAC |
| Zuza27a-targeting_spacer_F | ACCGCACTTTCAAGCTCTTCAAACTCTC CCCGGTGT |
| Zuza27a-targeting_spacer_R | AAACACACCGGGGAGAGTTTGAAGAGC TTGAAAGTG |
| Oligonucleotide sequences used for the construction of plasmid carrying dual-spacer mini-CRISPR array | |

| | |
|---|---|
| Olig_phiko_r_phifa_F | ACCGCCCTTGGGGAATGGCTTGAGGCGGCAACACGGGTTTCTCCCGTGCGCGCATGGAGGCACCGGTGAACCCGTCCAGACCATCGGCGAATACGAA |
| Olig_phiko_r_phifa_R | AAACTTCGTATTCGCCGATGGTCTGGACGGGTTCACCGGTGCCTCCATGCGCGCACGGGAGAAACCCGTGTTGCCGCCTCAAGCCATTCCCCAAGGG |
| Construction of plasmid encoding the LalCas12r(D238A) mutant | |
| cas12r_D238A_F | CTGGGGGTGGCCGTGAACGCCCG |
| cas12r_D238A_R | GCGTTCACGGCCACCCCCAGCAC |
| pMK18_Gib_HindIII_F | AAAACGACGGCCAGTGCCAAGCTTGCATGC |
| pMK18_Gib_EcoRI_R | ACAGCTATGACATGATTACGAATTCGAGCTCGGTAC |
| Construction of plasmids encoding LalCas12r and a mini-CRISPR array consisting of a single repeat and an 18-nt spacer targeting phage phiKo | |
| LalCas12r_put_promoter_SalI_F | CTGCAGGTCGACTTGACACTCAGGC |
| 18_bp_phiko_sp_EcoRI_R | ATTAAGAATTCCAAGCCATTCCCCAAGGGCGG |
| Colony PCR analysis | |
| cas12r_537R_F | GAGCGCCTAGGTCCGGATCG |
| ins_R | CACTTTATGCTTCCGGCTCGTATG |
| Analysis of phiKo phage escapers diversity | |
| phiKo_ps_F | AACGTACCCCTCCTCATGGA |
| phiKo_ps_R | CCGTGTGGTGCAGGACAATA |

**Supplementary Table S2.** Plasmids used in this study. All plasmid files are provided in supplementary data.

| **Plasmid name** | **Description** | **Figures/Supplementary Figures** |
|---|---|---|

| | | |
|---|---|---|
| pUC19_*lalcas12r*_mini-CRISPR (*lacZ*-alpha reporter region as a spacer) | Plasmid for insertion of spacers into the mini-CRISPR array | **-** |
| pMK18 | An empty vector was used as a control in plasmid transformation interference assays, escaper cell analysis, and identification of small RNAs associated with the LalCas12r–CRISPR system in *T. thermophilus* HB27c. | **3A,B, S9, S10A, S12B** |
| pMK18_*lalcas12r*–r_GT_r | A pMK18-derived plasmid encoding LalCas12r and a mini-CRISPR array with a genome-targeting spacer was used in plasmid transformation interference assays, and escaper cell analysis in *T. thermophilus* HB27c. | **3A,B, S9, S10A,B** |
| pMK18_*lalcas12r*–r_ST_r | A pMK18-derived plasmid encoding LalCas12r and a mini-CRISPR array with a self-targeting spacer was used in plasmid transformation interference assays, and escaper cell analysis in *T. thermophilus* HB27c. | **3A,B, S9, S10A,B** |
| pMK18_*lalcas12r*–r_NT_r | A pMK18-derived plasmid encoding LalCas12r and a mini-CRISPR array with a non-targeting spacer was used as a control in plasmid and phage transformation interference assays, and escaper phage analysis in *T. thermophilus* HB27c. | **3A,B, 4, S9, S11A** |
| pMK18_*dlalcas12r*–r_GT_r | A pMK18-derived plasmid encoding catalytically inactive LalCas12r(D238A) and a mini-CRISPR array with a genome-targeting spacer was used in plasmid transformation interference assays in *T. thermophilus* HB27c. | **3A,B** |

| | | |
|---|---|---|
| pMK18_*lalcas12r*–r_phiKo_r | A pMK18-derived plasmid encoding LalCas12r and a mini-CRISPR array with a phiKo phage-targeting spacer was used in phage transformation interference assays, and escaper phage analysis in *T. thermophilus* HB27c. | **4, S11** |
| pMK18_*lalcas12r*–r_phiFa_r | A pMK18-derived plasmid encoding LalCas12r and a mini-CRISPR array with a phiFa phage-targeting spacer was used in phage transformation interference assays | **4** |
| pMK18_*lalcas12r*–r_Zuza27_r | A pMK18-derived plasmid encoding LalCas12r and a mini-CRISPR array with a Zuza27 phage-targeting spacer was used in phage transformation interference assays | **4** |
| pMK18_*lalcas12r*–r_phiKo_r_phiFa_r | A pMK18-derived plasmid encoding LalCas12r and a dual-spacer mini-CRISPR array with phiKo and phiFa-targeting spacers was used in identification of small RNAs associated with the LalCas12r–CRISPR system in *T. thermophilus* HB27c. | **5A, S12A,B** |
| pMK18_*dlalcas12r*–r_phiKo_r_phiFa_r | A pMK18-derived plasmid encoding the catalytically inactive LalCas12r(D238A) and the dual-spacer mini-CRISPR array with phiKo and phiFa-targeting spacers was used in identification of small RNAs associated with the dLalCas12r–CRISPR system in *T. thermophilus* HB27c. | **5A, S12A,B** |
| pMK18_*lalcas12r*–r_phiKo(18nt 5' segment) | A pMK18-derived plasmid encoding the LalCas12r and a mini-CRISPR array consisting of one repeat and an 18 nt phiko-targeting | **5B, S12A,B** |

| | |
|---|---|
| | spacer was used in analysis of small RNAs associated with the LalCas12r–CRISPR system in *T. thermophilus* HB27c. |

**Supplementary Table S3.** Lalka27a genes classified according to the dynamics of transcript abundances.

| start | end | strand | gene id | product | time class |
|---|---|---|---|---|---|
| 27 | 404 | - | 1 | Ulx-like anti-restriction protein | late |
| 397 | 6621 | - | 2 | DarB-like anti-restriction protein | late |
| 6747 | 7469 | + | 3 | ssDNA binding protein | middle |
| 7659 | 8942 | - | 4 | hypothetical protein | late |
| 9045 | 9293 | + | 5 | hypothetical protein | pre-early |
| 9299 | 9967 | + | 6 | hypothetical protein | pre-early |
| 10005 | 10778 | - | 7 | hypothetical protein | late |
| 10789 | 11217 | - | 8 | hypothetical protein | late |
| 11214 | 11843 | - | 9 | hypothetical protein | late |
| 11895 | 13658 | + | 10 | portal protein | late |
| 13645 | 14595 | + | 11 | minor head protein | late |
| 14659 | 15639 | + | 12 | vWA domain-containing protein | middle |
| 15636 | 16982 | + | 13 | hypothetical protein | middle |
| 16979 | 18877 | + | 14 | AAA family ATPase | middle |
| 18880 | 19038 | + | 15 | hypothetical protein | middle |
| 19071 | 19295 | + | 16 | hypothetical protein | middle |
| 19360 | 21381 | + | 17 | terminase large subunit | late |
| 21378 | 21986 | - | 18 | hypothetical protein | middle |
| 21983 | 22372 | - | 19 | hypothetical protein | middle |
| 22314 | 23120 | - | 20 | dNMP kinase | middle |
| 23113 | 23745 | - | 21 | nucleotidase | middle |
| 23732 | 25003 | - | 22 | exonuclease | middle |
| 25143 | 25457 | - | 23 | hypothetical protein | late |
| 25457 | 25876 | - | 24 | hypothetical protein | late |
| 25939 | 26199 | - | 25 | hypothetical protein | late |
| 26220 | 26435 | - | 26 | hypothetical protein | late |
| 26438 | 27091 | - | 27 | M23 family peptidase | late |

| | | | | | |
|---|---|---|---|---|---|
| 27101 | 27550 | - | 28 | hypothetical protein | late |
| 27844 | 28260 | - | 29 | hypothetical protein | late |
| 28269 | 29333 | - | 30 | hypothetical protein | late |
| 29335 | 29721 | - | 31 | hypothetical protein | late |
| 29752 | 31038 | - | 32 | hypothetical protein | late |
| 31039 | 32100 | - | 33 | baseplate wedge subunit | late |
| 32097 | 32456 | - | 34 | baseplate protein | late |
| 32460 | 33029 | - | 35 | hypothetical protein | late |
| 33022 | 33648 | - | 36 | hypothetical protein | late |
| 33674 | 33937 | - | 37 | PAAR-domain spike tip protein | late |
| 33937 | 34371 | - | 38 | putative zinc metallopeptidase | late |
| 34402 | 35178 | - | 39 | baseplate protein | late |
| 35175 | 35387 | - | 40 | hypothetical protein | late |
| 35377 | 40524 | - | 41 | virionic helicase | late |
| 40588 | 41970 | - | 42 | tail protein | late |
| 41971 | 44640 | - | 43 | M23 family peptidase | late |
| 44625 | 44879 | - | 44 | hypothetical protein | late |
| 44845 | 45261 | - | 45 | tail assembly chaperone | late |
| 45332 | 45793 | - | 46 | virion structural protein | late |
| 45807 | 47546 | - | 47 | tail sheath | late |
| 47560 | 47769 | - | 48 | hypothetical protein | late |
| 47807 | 48679 | - | 49 | structural protein | late |
| 48689 | 49573 | - | 50 | hypothetical protein | late |
| 49570 | 50475 | - | 51 | hypothetical protein | late |
| 50476 | 51279 | - | 52 | tail fiber protein | late |
| 51309 | 51713 | - | 53 | yqbF-like protein | late |
| 51815 | 53509 | + | 54 | LalCas12r | middle |
| 53960 | 55468 | - | 57 | MCP | late |
| 55465 | 56112 | - | 58 | hypothetical protein | late |
| 56126 | 57046 | - | 59 | phage prohead protease | late |
| 57085 | 57999 | + | 60 | recombinase | late |
| 58009 | 58227 | + | 61 | hypothetical protein | late |
| 58174 | 59019 | - | 62 | DNMTa | middle |
| 59016 | 59861 | - | 63 | DNMT | middle |

| | | | | | |
|---|---|---|---|---|---|
| 59848 | 60042 | - | 64 | hypothetical protein | middle |
| 60008 | 60277 | - | 65 | hypothetical protein | middle |
| 60274 | 60897 | - | 66 | primase | middle |
| 60885 | 62183 | - | 67 | DnaB-like replicative helicase | middle |
| 62173 | 63108 | - | 68 | replication initiation O-like protein | middle |
| 63195 | 63326 | - | 69 | hypothetical protein | middle |
| 63443 | 63535 | + | 70 | hypothetical protein | late |
| 63641 | 64696 | - | 71 | DNA polymerase processivity factor | middle |
| 64771 | 65025 | - | 72 | hypothetical protein | middle |
| 65037 | 65819 | - | 73 | RecT-like ssDNA annealing protein | middle |
| 65845 | 66024 | - | 74 | hypothetical protein | middle |
| 66021 | 66344 | - | 75 | hypothetical protein | early |
| 66365 | 66547 | - | 76 | hypothetical protein | early |
| 66544 | 66825 | - | 77 | hypothetical protein | early |
| 66822 | 66977 | - | 78 | hypothetical protein | early |
| 66970 | 67593 | - | 79 | hypothetical protein | early |
| 67586 | 68011 | - | 80 | hypothetical protein | early |
| 68033 | 68437 | - | 81 | transcriptional regulator | early |
| 68552 | 69232 | + | 82 | LexA-like regulator | middle |
| 69234 | 69887 | + | 83 | hypothetical protein | middle |
| 69887 | 71077 | + | 84 | integrase | middle |

**Supplementary Table S4.** The list of Lalka27a virion proteins

| **№** | **Match to** | **Accession** | **Expect** | **Score** | **Nominal mass (Mr)** | **pI value** | **Number of mass values searched** | **Number of mass values matched** |
|---|---|---|---|---|---|---|---|---|
| 1 | dNMP kinase | XKB89803.1 | 2.4e-010 | 141 | 28911 | 07.08 | 39 | 12 |
| 2 | MCP | XKB89840.1 | 1.2e-023 | 275 | 54613 | 6.23 | 28 | 21 |
| 3 | DarB-like anti-restriction protein | XKB89785.1 | 1.2e-037 | 415 | 230251 | 6.20 | 88 | 62 |
| 4 | tail | XKB898 | 1.2e-018 | 225 | 60840 | 6.51 | 59 | 26 |

| | | | | | | | | |
|---|---|---|---|---|---|---|---|---|
| | sheath | 30.1 | | | | | | |
| 5 | hypothetical protein Lalka27a_4 | XKB89787.1 | 2,00E-09 | 133 | 47756 | 9.44 | 43 | 15 |
| 6 | putative helicase | XKB89824.1 | 3.9e-010 | 140 | 192212 | 6.66 | 115 | 39 |
| 7 | virion structural protein | XKB89829.1 | 7.8e-029 | 327 | 16178 | 4.91 | - | - |
| 8 | hypothetical protein Lalka27a_59 | XKB89842.1 | 9.8e-012 | 156 | 33845 | 10.10 | 26 | 12 |

**Supplementary Table S5.** Protospacers matching LalCas12r-CRISPR spacer and their genomic context. For details see **Fig. 2A** legend.

| Spacer origin and position in the CRISPR array | Spacer sequence | Spacer length, bp | Organism with protospacer | Protospacer sequence | Protospacer start coordinate | Protospacer coordinate | Protospacer genomic context | Spacer–protospacer matches (nt) | Spacer–protospacer identity (%) |
|---|---|---|---|---|---|---|---|---|---|
| Lalka27b_sp1 | GACGGGAGGGGGTACGAGTGGGAGGTGCCCCT | 32 | *T. scotoductus* Uz79 | GACGGGAGGGGGTACGAGTGGGAGGTGCCCCT | 148275 | 148306 | Integrative element | 32 | 100 |
| Lalka27b_sp3 | GAAATCGCCAAAGCGGGGCCCGCCGTCCTCCT | 32 | *T. scotoductus* Uz79 | GAAATCGCCAAAGCGGGGCCCGCCGTCCTCCT | 147264 | 147295 | Integrative element | 32 | 100 |
| Lalka27b_sp6; Lalka27d_sp2; Lalka27c_sp4 | TCTTGGTGGCGGGGTCATAGACGACCCACCC | 32 | *Thermus sp.* KM4338-28 | TCTTGGTGGCGGGCTCATAGACCACCCACCC | 10168 | 10138 | Putative conjugative plasmid (*) | 29 | 96.875 |
| Lalka27a_sp1 | GTCAGGGGAGGTGTCCCCGTTTTTGTCCCCGA | 32 | *Thermus sp.* H19.2 | GTCAGGGGAGGTGTCCCCGTTTTTGTCCCCGA | 1440505 | 1440536 | Integrative element | 32 | 100 |
| Lalka8_sp1 | TCTTACCCGAAATGCACCCCTCTACGGTCTAC | 33 | *T. thermophilus* TMY | TCTTGCCCGAAATGCACCCGTCTACGGTCTAC | 1013124 | 1013093 | Integrative element | 30 | 96.96969697 |
| Lalka8_sp2 | GTGGGGCGGGTTTGGCGCATCCCGAGGGCCGC | 32 | *T. antranikianii* isolate RBS10-92 | GTGGGCCGGGTTTGGCGCATCCCGAGGGCCGC | 1416301 | 1416270 | Integrative element | 31 | 100 |
| Lalka8_sp3_sp4 | ACCGCCCAGGCCCGGGCCGTCCACATCCTGACC | 33 | *T. thermophilus* SNM1-1 | ACCGCCCAGGCCCGGGCGGTGCACATCCTGACC | 610684 | 610716 | Integrative element (+) | 31 | 100 |
| Lalka8_sp5 | TCCGGGCGCGGTTTGAGTGGACGCTTTACAAG | 32 | *T. thermophilus* SNM6-6 | TCCGGGCGCGGTTTGAGTGGACGCTTTACAAG | 1089858 | 1089889 | Integrative element | 32 | 100 |
| Lalka8_sp7 | TAGTGCCGATGGACCTGGCACGCCCCGGCACGC | 33 | *T. thermophilus* SNM1-7 | TAGTGCCGATGGACCTGGCACGCCCCGGCACGC | 850799 | 850831 | Integrative element | 33 | 100 |
| Ga0181858100427_s | GTAAAAACGCTTCCCAATCCGC | 32 | *T. antranikianii* YIM 73052 | GTAAAAACGCTTCCCAATCCGC | 17415 | 17384 | Integrative element | 32 | 100 |

| | | | | | | | | | |
|---|---|---|---|---|---|---|---|---|---|
| p3 | ACCCCGTGCC | | | ACCCCGTGCC | | | | | |
| Ga0181858100427_sp4 | GAGCCGTAGGTGTGGCGGAGGTCGTGGACGCG | 32 | *T. antranikianii* YIM 73052 | GAGCCGTAGGTGTGGCGGAGGTCGTGGACGCG | 143879 | 28204 | Integrative element | 32 | 100 |
| Ga0181858100427_sp6 | GCGGTTCTGAAACGCGTTGGCATCGGCCAGGA | 32 | *T. thermophilus* AA1-1 | GCGGTTCTAAAACGCGTTGGCATCGGCCAGGA | 891503 | 891472 | Integrative element | 31 | 100 |
| ERZ4756919.5876_sp1 | GAGTTCTACCCCGAACGGCTCGGGGACGGG | 32 | *T. aquaticus* Y51MC23 | GAGCTCCACCCCGAAGGGCTCGGGGACGGG | 706764 | 706793 | Integrative element | 29 | 100 |
| ERZ4756919.5876_sp2 | CGGGGATCCACGTGCGGGTTGAGGGCCTACCCC | 33 | *T. thermophilus* SNM1-7 | CGGGGATCCACGTGCGGGTTGAGGGCCTACCCC | 846507 | 846475 | Integrative element | 33 | 100 |
| Lalka27c_sp1 | GAATGCCGTTCCCAACGGAAACTACGGGACAC | 32 | *Thermus sp.* H19.2 | GAATGCCGTTCCCAACGGAAACTACGGGACAC | 1439546 | 1439515 | Integrative element | 32 | 100 |
| Lalka27c_sp2 | TTGTCTCTTGCGCGAAACCGTGCCATGCCGC | 32 | *Thermus sp.* H19.2 | TTGCCTCTTGCGCGAAACCGTGCCATGCCGC | 1440041 | 1440071 | Integrative element | 30 | 96.875 |
| Tsp_93170_prophage_sp1 | CCCCGGCAGGTGGTGGCCGTCCCCTACAGCA | 31 | *T. aquaticus* Y51MC23 | CCCCGGCAGGTGGTGGCCGTCCCCTACAGCA | 1046770 | 1046800 | Prophage | 31 | 100 |
| Tsp_93170_prophage_sp2 | GTTGGCCGCCAATGGTCGTACGAGCCAGCCGT | 32 | *T. scotoductus* 38_S38 | GTTGGCCGCCAATGGTCGTACGAGCCAGCCGT | 1324 | 1293 | Putative plasmid | 32 | 100 |

**Supplementary Table S6.** Protospacer adjacent sequences containing putative PAMs.

| Flank ID | Spacer origin and position in the CRISPR array | 8 bp 5' protospacer-adjacent sequence |
|---|---|---|
| flank 1 | Lalka8_sp7 | GGGGGAGG |
| flank 2 | Lalka8_sp5 | GGAGAAAG |
| flank 3 | Lalka27b_sp1 | ACCTGAAG |

**Supplementary Table S7.** CFU counts upon transformation of *T. thermophilus* HB27c cells with distinct *lalcas12r*-CRISPR-bearing constructs**.**

| plasmid | replica | count |
|---|---|---|
| pMK18 | 1 | 482 |
| pMK18 | 2 | 1078 |
| pMK18 | 3 | 568 |
| pMK18_*lalcas12r*-r_NT_r | 1 | 2526 |
| pMK18_*lalcas12r*-r_NT_r | 2 | 1729 |
| pMK18_*lalcas12r*-r_NT_r | 3 | 1900 |
| pMK18_*dlalcas12r*-r_GT_r | 1 | 2207 |
| pMK18_*dlalcas12r*-r_GT_r | 2 | 1319 |
| pMK18_*dlalcas12r*-r_GT_r | 3 | 1820 |
| pMK18_*lalcas12r*-r_GT_r | 1 | 112 |

| | | |
|---|---|---|
| pMK18_*lalcas12r*-r_GT_r | 2 | 327 |
| pMK18_*lalcas12r*-r_GT_r | 3 | 395 |
| pMK18_*lalcas12r*_r_ST_r | 1 | 317 |
| pMK18_*lalcas12r*_r_ST_r | 2 | 326 |
| pMK18_*lalcas12r*_r_ST_r | 3 | 3 |

**Supplementary Table S8.** Description of the TnpB branch containing LalCas12r and LlkCas12r.

| Tree Id | Source organism | Accession | Thermal type |
|---|---|---|---|
| 16749 | *Hydrogenivirga okinawensis* JCM 13302 | BGR20498.1 | thermophilic |
| 10598 | *Carboxydothermus ferrireducens* DSM 11255 | NYE57029.1 | thermophilic |
| 22408 | *Caldicellulosiruptor morganii* DSM 8990 | WP_045170344.1 | thermophilic |
| 1506 | *Caldicellulosiruptor acetigenus* I77R1B | WP_013432624.1 | thermophilic |
| 18209 | *Caldicellulosiruptor bescii* DSM 6725 | WP_015907403.1 | thermophilic |
| LlkCas12r | metagenomic contig (Ga0079992_140590) | Ga0079992_1405904 | thermophilic |
| LalCas12r | Lalka8 | XKB89754.1 | thermophilic |
| LalCas12r | Lalka27a | XKB89837.1 | thermophilic |
| LalCas12r | Lalka27b | XKB89922.1 | thermophilic |
| LalCas12r | Lalka27c | pending | thermophilic |
| LalCas12r | Lalka27d | pending | thermophilic |
| 10879 | *Thermus aquaticus* | WP_424155722.1 | thermophilic |
| 4372 | *Thermus thermophilus* TMY | BAW02830.1 | thermophilic |
| 10970 | *Thermocrinis sp.* | WP_299197249.1 | thermophilic |
| 10547 | *Thermocrinis sp.* | WP_424182520.1 | thermophilic |
| 12726 | *Thermocrinis sp.* | MCC6063678.1 | thermophilic |
| 8842 | *Thermoflexus sp.* | MDT7949546.1 | thermophilic |
| 4793 | *Thermoflexus sp.* | MDT7947265.1 | thermophilic |
| 22132 | *Sulfurihydrogenibium sp.* YO3AOP1 | WP_012460240.1 | thermophilic |

| | | | |
|---|---|---|---|
| 17825 | *Hydrogenobaculum sp.* SHO | WP_015419257.1 | thermophilic |
| 15808 | *Candidatus Caldipriscus sp.* | MCC6011663.1 | thermophilic |
| 1528 | *Thermocrinis ruber* DSM 23557 | WP_051402152.1 | thermophilic |
